# Multi-omic dissection of RNA control reveals convergent *cis*-regulatory programs during glucose starvation in yeast

**DOI:** 10.64898/2026.07.20.739609

**Authors:** Orlane Rossini, Shafi Mahmud, Nikolay Shirokikh, Alice Cleynen

**Author notes:** Corresponding authors: Nikolay Shirokikh, Alice Cleynen.

## Abstract

Cells reprogram gene expression at multiple, often independently studied, regulatory layers during nutrient stress. Here we use direct RNA sequencing of polysome-fractionated transcripts that simultaneously measures transcript abundance, mRNA degradation, poly(A) tail length, and three RNA modifications (m5C, m6A, pseudouridine) to build an integrated, multi-omic view of the *Saccharomyces cerevisiae* response to acute glucose starvation. Lasso-penalised mixture-of-regressions clustering of stochastic translation efficiency (δSTE) against this combined feature set partitions 5,033 transcripts into five translational programs with distinct functional identities, from a strongly upregulated translation-machinery cluster to a strongly repressed glycosylation cluster. CDS sequence composition discriminates these extremes, while 5′ and 3′UTR composition contributes only marginal, largely non-significant effects after length correction. Extending this analysis, we show that the repressed glycosylation program is distinguished by a convergent stack of independent *cis*-regulatory signatures: weak Kozak initiation context, high upstream open reading frame burden, excess RNA secondary structure specifically over the start codon, and depletion for the targets of major stabilising and export-associated RNA-binding proteins. In other programmes, translation efficiency tunes with initiation-context strength indicating that downstream regulatory layers coregulate translation. These results provide an integrated, feature-resolved map of post-transcriptional regulation during acute nutrient stress.

## 1. Introduction

The ability to sense and adapt to a fluctuating environment is a fundamental property of living cells, and among the best-characterised adaptive responses is that triggered by nutrient limitation, which forces cells to rapidly reprogram gene expression, metabolism, and protein synthesis. The budding yeast *Saccharomyces cerevisiae* has long served as a powerful model for dissecting such responses, owing to its genetic tractability, the conservation of core molecular pathways with higher eukaryotes, and its comprehensive genomic resources. In their natural habitat, yeast cells encounter diverse stresses, including heat shock, oxidative, osmotic and heavy-metal stress, and DNA damage, that elicit overlapping yet distinct transcriptional and post-transcriptional programs(1,2). A particularly instructive paradigm is glucose limitation and starvation, which drives a profound physiological transition: cells rapidly exit proliferation, remodel their proteome, and enter a quasi-quiescent state that permits long-term survival(3,4). This transition is not a passive shutdown but an active, coordinated reprogramming that reallocates cellular resources, most notably the translational machinery, from growth-promoting functions toward metabolic and stress-protective programs.

The transcriptional response to glucose starvation has been characterised extensively. Landmark genomic surveys established that hundreds of transcripts change in abundance across diverse stresses, defining a core “environmental stress response” (ESR) that is induced or repressed in a stereotyped manner largely independent of the specific stress(1). Under glucose limitation, transcripts encoding ribosomal proteins and translation factors are dramatically downregulated, while those encoding enzymes of gluconeogenesis, the glyoxylate cycle, fatty acid oxidation, and stress-protective functions are upregulated(5,6). Beyond steady-state transcript abundance, it has become clear that the control of translation plays an equally critical role in shaping the cellular response. A canonical example is the translational induction of GCN4, encoding a transcriptional activator of amino acid biosynthesis genes: its expression is controlled by upstream open reading frames in its 5ʹUTR and is rapidly induced under glucose or amino acid limitation, demonstrating how fast, specific translational changes can act as an initial stress response before triggering longer-term transcriptional reprogramming(7,8). More broadly, polysome profiling and ribosome footprinting studies have revealed that the relationship between mRNA levels and ribosome occupancy is far from deterministic: many transcripts maintain or even increase their translational efficiency under stress conditions despite declining abundance, while others are selectively excluded from the translational pool(9–12). This selective translational control allows cells to rapidly alter protein output without necessarily modifying transcription, and the partitioning of mRNAs between polysome-associated (actively translated) and ribosome-free fractions has emerged as a key regulatory event during glucose starvation, with distinct functional classes of transcripts showing differential engagement with the translational machinery(8). Yet accurately quantifying translational efficiency transcriptome-wide has remained technically challenging, since classical normalisation of ribosome footprints to total mRNA levels conflates translational control with changes in mRNA abundance and availability(12). To address this limitation, we recently developed enhanced translation complex profile sequencing (eTCP-seq) based on rapid *in vivo* crosslinking, together with an AI-based machine learning framework that computes a new, self-normalised measure of translational output, Stochastic Translation Efficiency (STE), which maps directly to absolute protein biosynthesis rates and is independent of mRNA abundance fluctuations(13). Applied to yeast undergoing 10-minute glucose depletion, STE revealed rapid translational upregulation of oxidative stress and sugar metabolism genes, illustrating the power of the approach to capture the earliest stages of stress-responsive translational reprogramming(13).

Yet the picture emerging from studies of transcript abundance and translational efficiency is almost certainly incomplete. It is now well established that mRNA molecules carry multiple molecular features that independently influence their translational fate, and that each of these can be dynamically remodelled under stress(14). mRNA stability is a primary determinant of transcript availability: degradation rates can change rapidly and independently of transcription rates, and shifts in mRNA half-life have been shown to account for a substantial fraction of the expression changes observed under stress(15,16). The length of the 3ʹ poly(A) tail constitutes another distinct regulatory axis: cytoplasmic deadenylation and re-polyadenylation dynamically tune translational competence and stability in a transcript-specific manner, with poly(A) tail length linked to ribosome recruitment and mRNA fate decisions(17–19).

Beyond the larger structural features, an expanding repertoire of chemical modifications of the RNA bases, collectively termed the epitranscriptome, has emerged as a further layer of translational regulation(14). N6-methyladenosine (m6A) influences translation efficiency, mRNA stability, and stress granule partitioning, and has been shown to be dynamically regulated under stress in mammalian systems and linked to mRNA fate decisions in yeast(20–23). 5-methylcytidine (m5C) in mRNA has been linked to translational control and mRNA nuclear export under specific cellular conditions(24–26). Pseudouridine (Ψ), the most abundant RNA modification, is dynamically induced in mRNAs under nutrient deprivation and heat shock in yeast, with direct consequences for decoding and translational fidelity; it is also deposited co-transcriptionally in pre-mRNA, affecting splicing and polyadenylation site choice(21,27,28).

Crucially, each of the regulatory axes, the stability, poly(A) length, and RNA modifications, has been studied predominantly in isolation and linked to translation individually. Whether and how these axes act in concert on overlapping or distinct transcript populations to collectively shape translational output during stress remains an open and fundamental question, and answering it requires measuring them together rather than one layer at a time.

Here we address this question through a simultaneous multi-omic characterisation of the yeast response to glucose starvation. Direct RNA sequencing of native transcripts is central to this strategy: because abundance, stability, poly(A) tail length and multiple base modifications can be read from the same individual molecules, these regulatory layers can be related to one another within a single biological sample rather than reconstructed from separately generated datasets. We first systematically assess how each regulatory axis (transcript abundance, mRNA stability, poly(A) tail length, m5C, m6A, and pseudouridine modifications) is remodelled under stress, in both light polysome-associated (up to 3 ribosomes, ribosome-disome-trisome, RDT) and heavy polysome-associated (more than 4 ribosomes, PS) fractions, connecting each feature to translational engagement where possible. We then apply the STE framework(13) to obtain a direct readout of translational output, and ask to what extent the individual and combined patterns of these regulatory features can account for the translational landscape observed during glucose starvation. Rather than treating the layers as parallel, independent readouts, we model them jointly and find that their relationship to translational output is non-additive: individual features predict opposite outcomes in different transcript populations, and coherent regulation becomes apparent only once transcripts are resolved into distinct translational programs (**Figure 1**). This integrated analysis identifies classes of transcripts subject to coordinated, multi-feature control and enriched for critical metabolic and translational functions, as well as it suggests that such transcripts are defined, at the sequence level, for particular translational fates that starvation then enacts.

**Figure 1.**
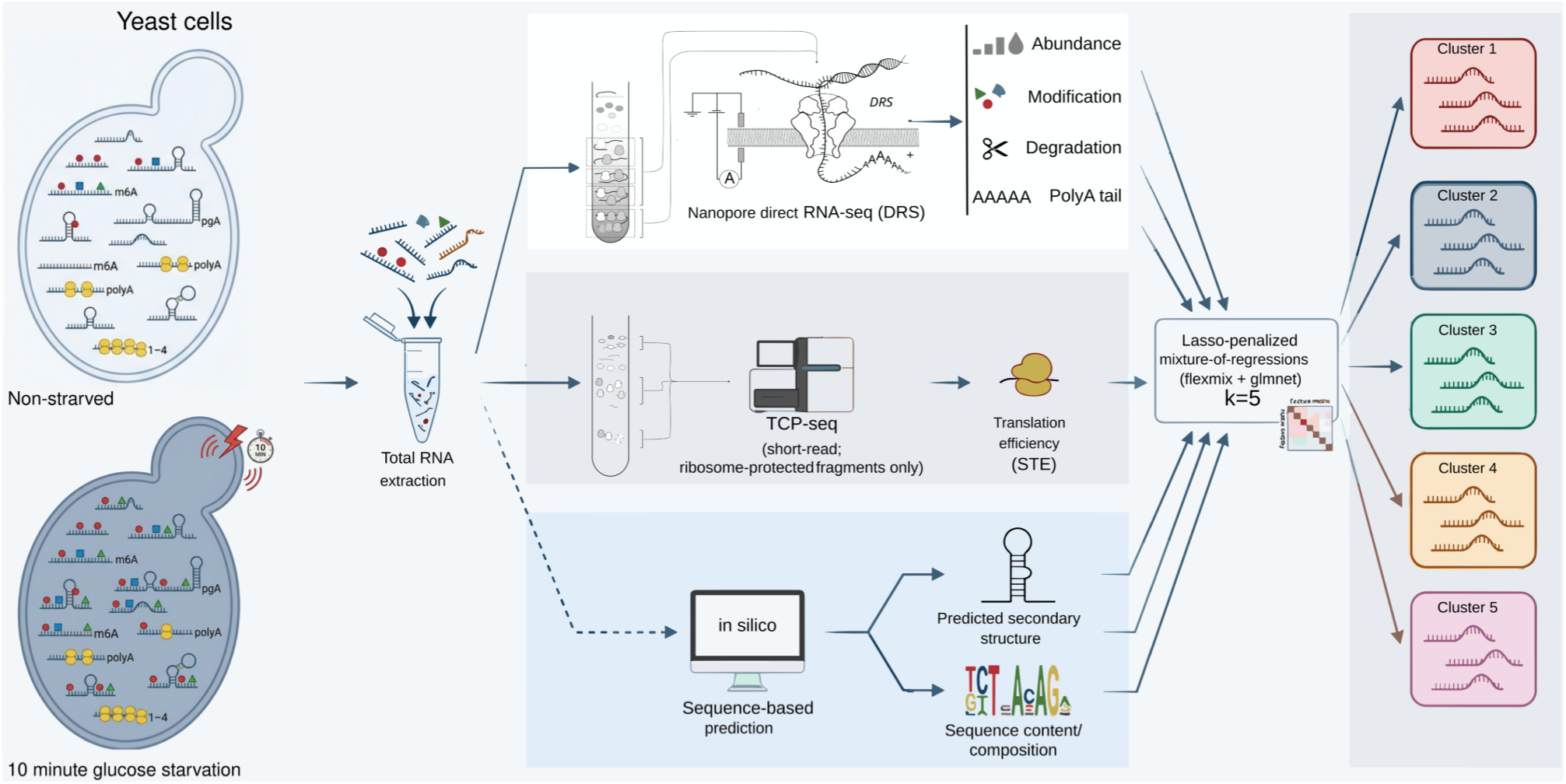
Multi-omic experimental and computational workflow for identifying stress-responsive translational programs in budding yeast. *Saccharomyces cerevisiae* cultures were profiled under two conditions, non-starved and after 10 minutes of acute glucose starvation, to capture the immediate translational stress response. Total RNA was extracted from each condition and characterised through three complementary analytical branches. (1) Nanopore direct RNA sequencing (DRS) on two polysome fractions provided native, single-molecule readout of transcript abundance, RNA base modifications, mRNA degradation, and poly(A) tail length. (2) TCP-seq of ribosome-protected fragments (short-read) yielded per-transcript estimates of translation efficiency (STE) across ribosome-loading fractions. (3) In silico sequence-based prediction supplied complementary structural and compositional features, namely predicted RNA secondary structure and sequence/codon composition. These feature layers were integrated in a lasso-penalised mixture-of-regressions model (flexmix + glmnet, k = 5), which groups transcripts according to the combination of features that best predicts the starvation-induced change in translation efficiency, yielding five transcript clusters (Clusters 1–5) with distinct regulatory signatures that are characterised throughout the remainder of the manuscript.

## 3. Results

### 3.1. Uni-omics investigation of response to stress

#### 3.1.1. Differential transcript representation in fractions

To characterise how nutrient availability influences translational engagement, we performed differential expression analysis using DESeq2 on both heavy polysome (PS) and light polysome (RDT) fractions, comparing starved and non-starved conditions. By intersecting the sets of transcripts significantly altered in both fractions, we identified 391 transcripts upregulated (occupied by the ribosomes more) in the non-starved condition and 323 transcripts upregulated in the starved condition (adjusted *p* < 0.01, abs(log_2_FC)>1).

To explore the relationship between differential expression and translation efficiency, we compared the average log_2_ fold change of each transcript between conditions with the corresponding difference in stochastic translation efficiency (STE) values (**Figure 2A**). This analysis revealed a striking divergence in translational strategy: transcripts upregulated in the starved condition (S10) generally exhibited higher STE values in S10, suggesting coordinated stabilisation and enhanced translation under nutrient stress. Conversely, transcripts upregulated in the non-starved condition showed a trend toward lower STE values in the same condition, indicating that despite higher abundance, their translational efficiency may be downregulated, possibly reflecting saturation of the translational machinery or regulatory buffering mechanisms. These findings suggest that nutrient status not only reshapes the translatome composition but also modulates the efficiency with which transcripts are translated, highlighting a complex interplay between transcript abundance and translational control in adaptive cellular responses.

**Figure 2.**
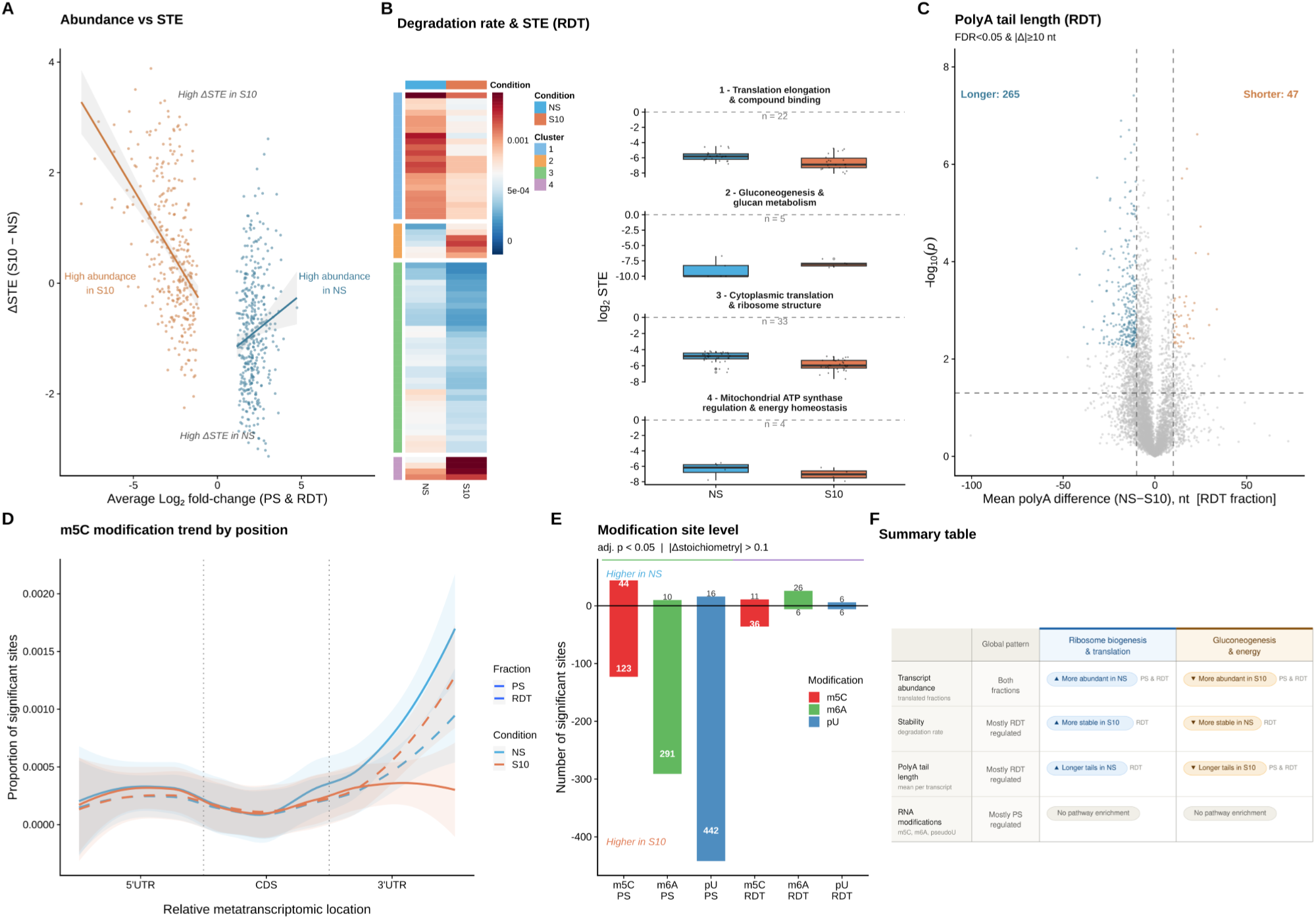
Multi-layer post-transcriptional regulation during nitrogen and glucose starvation. **(A)** Relationship between transcript abundance changes and stochastic translation efficiency (STE) in the heavy polysome (PS) fraction. Each point represents one transcript; x-axis shows the average log₂ fold-change in abundance (NS vs S10), y-axis shows δSTE (S10 − NS). Coloured points represent the quadrants of interest: orange for higher abundance in S10; blue for higher abundance in NS. Linear regression lines are shown per condition. **(B)** Differential mRNA stability and its relationship to STE. Left: Heatmap of degradation rates in the RDT fraction for transcripts in four stability-based clusters (colour bar at right). Rows are genes; columns are NS and S10. Right: Per-cluster log₂ STE distributions (NS vs S10) shown as vertical boxplots. Cluster annotations indicate the dominant GO biological process enrichment. **(C)** Volcano plot of polyA tail length changes between NS and S10 in the RDT fraction. Each point represents one transcript; x-axis shows mean polyA length difference (NS − S10, nt); y-axis shows −log₁₀(p-value). Significant transcripts (FDR < 0.05 and |Δ| ≥ 10 nt) are coloured blue (longer in NS) or red (shorter in NS). Counts of significant transcripts are shown. **(D)** m5C modification rate across the relative metatranscriptomic location (5’UTR, CDS, 3’UTR). Loess smoothed curves are shown for all four condition × fraction combinations: NS PS (solid blue), S10 PS (solid orange), NS RDT (dashed blue), S10 RDT (dashed orange). Shaded ribbons indicate 95% confidence intervals. Dashed vertical lines indicate UTR/CDS boundaries. **(E)** Number of significant RNA modification sites per comparison. Each bar represents one modification type (m5C red, m6A green, pU blue) in one fraction (PS or RDT). Bars above zero indicate sites with higher stoichiometry in NS; bars below zero indicate higher stoichiometry in S10. Numbers show absolute site counts. Significance thresholds: BH-adjusted p < 0.05, |Δstoichiometry| > 0.1. **(F)** Summary matrix of uni-omic regulatory patterns. Rows: four regulatory layers. Columns: global pattern and two functionally characterised gene groups (ribosome biogenesis & translation; gluconeogenesis & energy). Coloured badges indicate the direction of regulation (blue ▲ = elevated in NS; amber ▼ = elevated in S10); subscript labels indicate the predominant fraction (PS or RDT). Grey pills indicate no significant pathway enrichment.

#### 3.1.2. Differential transcript stability

To investigate mRNA degradation dynamics under varying nutritional states, we quantified transcript-specific degradation rates in both starved and non-starved conditions across the two polysome fractions (RDT and PS) using the INDEGRA software. While very few hits had changing stability in the PS fraction, hierarchical clustering of degradation profiles in RDT fraction identified four distinct clusters exhibiting differential stability patterns between conditions (**Figure 2B**). Clusters 1 and 3, comprising transcripts more stable under starvation (S10), showed striking enrichment for translation-related functions. Cluster 1 was particularly enriched for translation elongation factors and anion binding proteins, while Cluster 3 demonstrated exceptional enrichment for cytoplasmic translation machinery and structural constituents of ribosomes. This coordinated stabilisation and abundance reduction of the translational apparatus under nutrient stress suggests a protective mechanism to maintain protein synthesis capacity during adverse conditions. In contrast, Clusters 2 and 4, containing transcripts more stable in the non-starved (NS) condition, did not yield significant gene ontology hits but notably included transcripts encoding enzymes involved in gluconeogenesis, glucanotransferase activity, pseudohyphal growth, ATP-dependent gluconeogenesis, and phosphatase regulation, processes associated with metabolic flexibility and stress adaptation.

To explore the relationship between transcript stability and translational efficiency, we examined STE values across the degradation-defined clusters (**Figure 2B**). This analysis revealed complex, cluster-specific coordination between stability and translation that reflects distinct regulatory strategies. Cluster 1 transcripts, while stabilised under starvation, exhibited significantly reduced translation efficiency in S10 compared to NS, suggesting a decoupling of stability from active translational engagement where these transcripts are preserved but not utilised. Cluster 3 transcripts, already stable under non-starved conditions and further stabilised in S10, maintained high translation rates across both conditions with only modest reduction under starvation, indicating sustained engagement of core ribosomal components despite nutrient stress. In contrast, Cluster 4 transcripts showed dramatic destabilisation in S10 with minimal corresponding changes in STE, suggesting that their removal reflects a wholesale downregulation rather than translational remodeling. Cluster 2 transcripts exhibited both very low translation efficiency in both conditions and increased degradation under starvation, indicating these poorly translated transcripts are selectively cleared during nutrient stress. These findings reveal that cells employ multiple mechanisms to reshape the translatome: core translational machinery is buffered against degradation, translation elongation factors are stabilised but kept translationally silent, while metabolic transcripts with low translational engagement are preferentially degraded, allowing rapid reallocation of cellular resources in response to changing nutrient availability.

#### 3.1.3. Differential 3ʹ poly(A) length

To examine how nutrient availability influences post-transcriptional regulation through poly(A) tail length modulation, we analysed poly(A) tail distributions across both PS and RDT fractions under non-starved (NS) and starved (S10) conditions. Global analysis revealed that the starved condition consistently exhibited longer poly(A) tails compared to the non-starved fraction, a trend that was particularly pronounced in the RDT fraction (t-test p-value 10^−27^ for PS, 10^−116^ for RDT; **Figure 3A**), suggesting that nutrient stress promotes accumulation of polyadenylated transcripts outside the active translational pool. Differential analysis at the transcript level identified 265 transcripts with significantly longer poly(A) tails and 47 with shorter tails in S10 RDT compared to NS RDT, while the PS fraction showed a more modest response with 62 transcripts displaying longer tails and 32 showing shorter tails in S10 *versus* NS (**Figure 2C**; **Figure 3B**). Gene ontology enrichment analysis of transcripts with extended poly(A) tails revealed a striking shift in functional priorities between conditions: in the non-starved state, polyadenylation preferentially targeted transcripts encoding ribosomal components and nucleolar proteins, functions associated with active growth and protein synthesis (**Figure 3C and 3E**). In contrast, under starvation, transcripts with longer poly(A) tails were enriched for oxidoreductase activity, generation of precursor metabolites and energy, glyoxylate metabolic processes, carbohydrate metabolic processes, and mitochondrial localisation, reflecting a metabolic reprogramming toward energy production and stress adaptation (**Figure 3D and 3F**). These findings indicate that poly(A) tail extension serves as a dynamic regulatory mechanism that selectively stabilises functionally coherent transcript sets in response to nutrient availability, with starvation triggering a systematic shift from growth-promoting to metabolic survival programs through coordinated polyadenylation of stress-adaptive transcripts.

**Figure 3.**
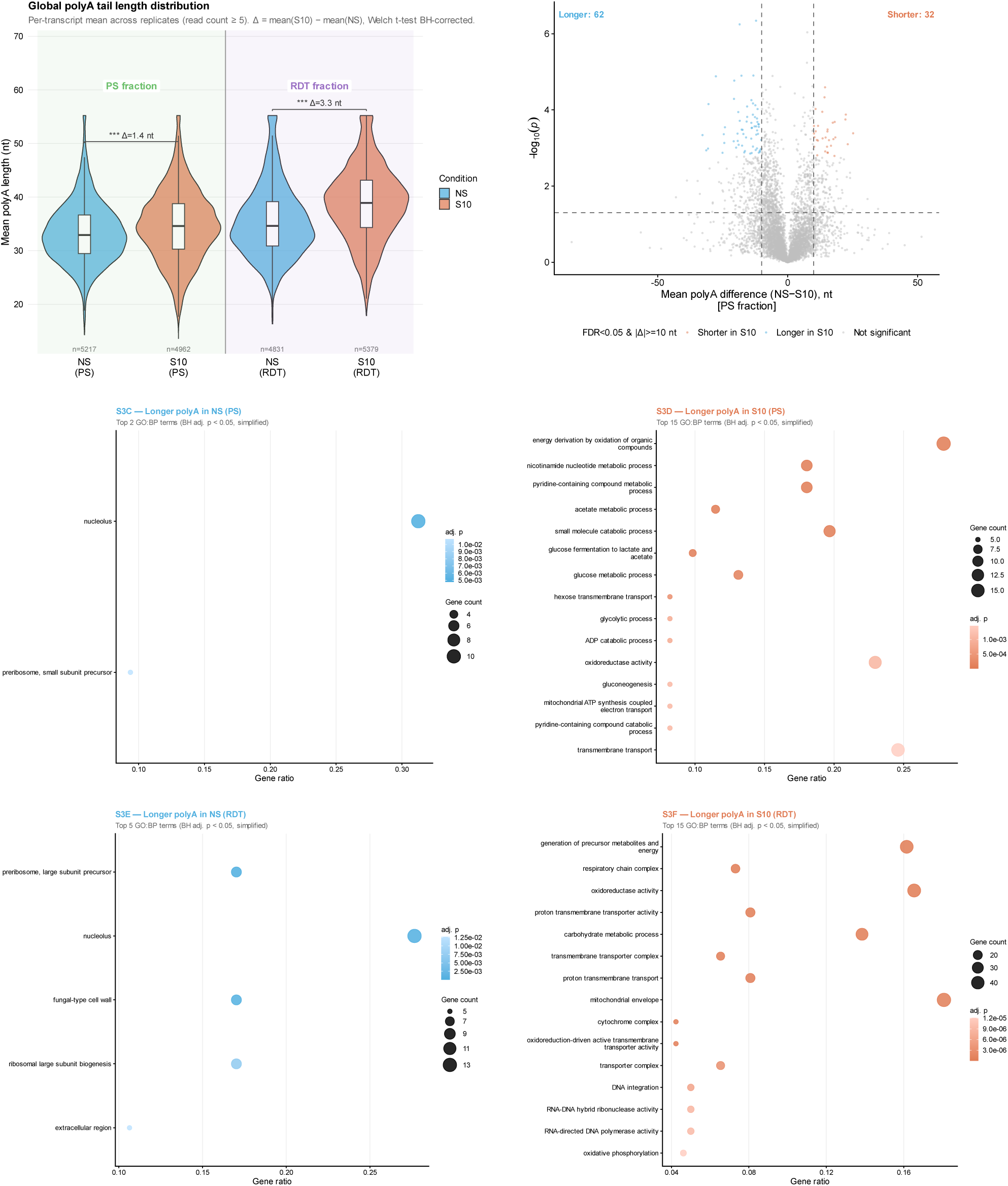
Global polyA tail length changes and pathway enrichment. **(A)** Global polyA tail length distributions by condition and fraction. Violin plots show per-transcript mean polyA length (read count ≥ 5, averaged across replicates) for all four condition × fraction groups. Embedded boxplots show median and IQR. Light background shading distinguishes PS (green) and RDT (purple) fractions, separated by a vertical line. n per group shown below each violin. Brackets show Welch t-test significance (BH-corrected) comparing NS vs S10 within each fraction; Δ = mean(S10) − mean(NS). **(B)** Volcano plot of polyA tail length changes in the PS fraction. Each point represents one transcript. Significant transcripts (FDR < 0.05, |Δ| ≥ 10 nt) are coloured blue (longer in NS) or orange (longer in S10). Counts of significant transcripts shown. **(C-D)** GO:BP enrichment for transcripts with longer polyA tails in NS (C) and S10 (D) in the PS fraction. Top GO terms shown (BH adj. p < 0.05, simplified). Point size = gene count; colour = adjusted p-value. **(E-F)** GO:BP enrichment for transcripts with longer polyA tails in NS (E) and S10 (F) in the RDT fraction.

#### 3.1.4. Differential RNA modifications

To investigate how nutrient availability influences epitranscriptomic regulation, we analysed three major RNA modifications, 5-methylcytidine (m5C), N6-methyladenosine (m6A), and pseudouridine (pU), across both fractions under non-starved (NS) and starved (S10) conditions. Metagene analysis revealed a modification-specific regional effect: m5C exhibited a significant reduction in modification rate within the 3ʹuntranslated region specifically in the S10 PS fraction (**Figure 2D**), while m6A and pU showed no consistent regional changes across conditions, albeit an overall reduction in modification rate in the starved condition in the PS fraction. At the individual site level, the PS fraction displayed extensive modification remodelling under starvation (**Figure 2E**), with 123 m5C sites, 291 m6A sites, and 442 pU sites showing increased stoichiometry in S10 compared to NS (*versus* only 44, 10, and 16 sites showing decreased stoichiometry, respectively). In contrast, the RDT fraction exhibited remarkable stability at the site level, with only 36 m5C, 6 m6A, and 6 pU sites showing increased stoichiometry (and 11, 26, and 6 showing decreases). Strikingly, however, when examining global stoichiometry shifts aggregated across all sites within individual transcripts, the RDT fraction showed more transcripts with overall modification changes than the PS fraction, despite fewer individual sites being altered.

Gene ontology enrichment analysis of transcripts with altered modification stoichiometry revealed a striking convergence on retrotransposon biology across all three modifications in the actively translating PS fraction under starvation. Transcripts with increased m5C, m6A, and pU stoichiometry in the S10 PS fraction were each independently enriched for terms related to retrotransposon activity, including RNA-directed DNA polymerase activity, RNA-DNA hybrid ribonuclease activity, DNA integration, retrotransposition, and retrotransposon nucleocapsid, reflecting the well-established activation of yeast Ty retrotransposons under nutrient stress conditions. Additional enrichment in cytoplasmic ribonucleoprotein granules and stress granule terms for m6A and pU further suggests coordinated remodelling of RNP assembly during starvation. The convergence of three chemically distinct modifications on the same functional gene class points to a layered epitranscriptomic response specifically targeting Ty element transcripts in translated mRNA pools. In contrast, the non-starved PS fraction showed a more metabolically specific signature: increased m5C in NS PS was enriched for polysaccharide and glycogen catabolic processes, trehalose metabolism, and energy reserve metabolic processes, consistent with the active carbon metabolism of exponentially growing cells.

In the RDT fraction, modification changes were largely below the threshold for GO enrichment, reflecting the smaller gene sets identified in this fraction. The sole exception was increased m5C stoichiometry in S10 RDT, which showed limited enrichment for phosphatidylinositol-4-phosphate binding alongside the retrotransposon signature, suggesting that this modification remodelling extends partially to non-polysome-associated transcripts under starvation.

#### 3.1.5 Multi-regulatory changes of metabolic and translational pathways

Having characterised individual regulatory layers (abundance, degradation, poly(A) tail length, and RNA modifications) in response to starvation, we next investigated whether these mechanisms act independently or converge on specific transcripts. Our pathway enrichment analyses revealed two major functional groups showing coordinated regulation: ribosomal/translational machinery and gluconeogenesis/energy metabolism. Ribosomal genes were consistently more abundant in non-starved conditions, more stable under starvation, and possessed longer poly(A) tails in the non-starved state. Conversely, gluconeogenesis genes showed the opposite pattern, with higher abundance under starvation, greater stability in non-starved conditions, and longer poly(A) tails under nutrient stress (**Figure 2F**). This suggested that stress-responsive genes may be subject to multi-layered regulatory control.

To systematically identify transcripts undergoing changes across multiple regulatory dimensions, we performed an intersection analysis of genes showing significant changes in abundance, degradation, poly(A) length, or modification status. Among the heavy polysome transcripts, we identified 1,279 genes with changes in at least one regulatory layer (**Figure 4A**). Strikingly, these regulatory changes were mostly exclusive of one another, however genes subject to multiple regulatory changes including degradation or poly(A) length were most often associated with changes in abundance or modification status. Notably, we identified genes with changes across three regulatory dimensions, representing transcripts under particularly complex regulatory control.

**Figure 4.**
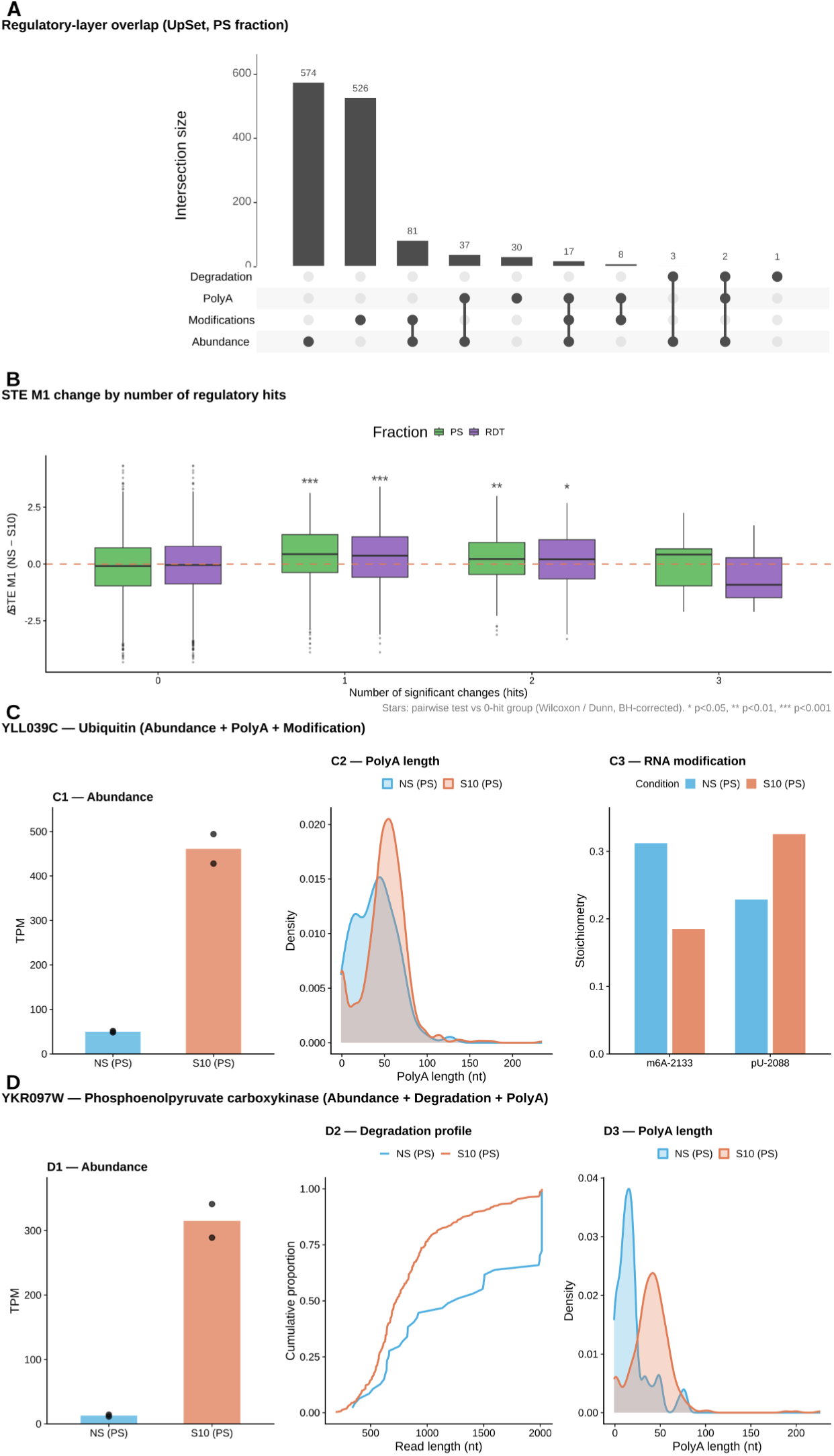
Multi-regulatory exclusivity patterns and relationship to translational reprogramming. **(A)** UpSet plot of multi-regulatory overlaps in the PS fraction. Bars show the number of transcripts with each combination of regulatory changes across four layers (transcript abundance, degradation, RNA modifications, polyA tail length). Horizontal bars at left show the total size of each regulatory set. Numbers above intersection bars indicate gene counts. **(B)** δSTE M1 (NS − S10) as a function of the number of significant regulatory changes (hits) per transcript, shown separately for the PS (green) and RDT (purple) fractions. Boxes show median and interquartile range; whiskers extend to 1.5×IQR. Outliers shown as individual points. Dashed orange line at y = 0. Statistics: Kruskal-Wallis test followed by pairwise Wilcoxon tests against the 0-hit group (BH-corrected); significance stars shown above each group. **(C)** Multi-regulatory example: YLL039C (ubiquitin; Abundance + PolyA + Modification, PS fraction). C1: TPM bar plot with replicate points (NS vs S10). C2: PolyA length density distribution. C3: RNA modification stoichiometry at two sites (m6A-2133, pU-2088) in NS and S10. **(D)** Multi-regulatory example: YKR097W (phosphoenolpyruvate carboxykinase; Abundance + Degradation + PolyA, PS fraction). D1: TPM bar plot with replicate points. D2: Cumulative read length distribution (degradation proxy). D3: PolyA length density distribution. In all panels, blue = NS, orange = S10.3.2. Multi-omics integration and mechanisms of response.

We next examined whether the number of regulatory changes correlated with translation efficiency. Focusing on STE model1, we stratified genes by their total number of significant regulatory hits (**Figure 4B**). Genes with multiple regulatory changes exhibited significantly higher efficiency compared to genes with fewer or no changes, particularly in the heavy polysome fraction (p-value <10^−15^ for 1 hit, <10^−2^ for 2 hits, not significant for 3 hits due to very small sample size).

To illustrate the biological relevance of multi-hit regulation, we examined two representative examples with changes across three regulatory layers. YLL039C, encoding ubiquitin, is essential for cellular stress response and protein quality control. Under starvation, YLL039C showed increased abundance, elevated m6A modification at position 2,133 in non-starved conditions, increased pseudouridine modification at position 2088 during starvation, and modestly longer poly(A) tails under stress (**Figure 4C**). This multi-layered control likely ensures precise ubiquitin levels during the transition to nutrient limitation, when protein turnover and quality control become critical.

YKR097W, encoding phosphoenolpyruvate carboxykinase (PCK1), represents a key regulatory node in gluconeogenesis. This gene exemplified the gluconeogenesis regulatory signature: dramatically increased abundance under starvation (20-fold by TPM), with a shift towards less stable transcripts in non-starved conditions, and substantially longer poly(A) tails during nutrient stress (**Figure 4D**). The cumulative degradation profile revealed distinct read length distributions between conditions, consistent with altered mRNA stability. The coordinated regulation of YKR097W across multiple layers likely enables the rapid metabolic switch from glycolysis to gluconeogenesis upon nutrient depletion.

Together, these results demonstrate that stress-responsive genes are not simply up- or down-regulated, but rather undergo coordinated changes across transcriptional, post-transcriptional, and epitranscriptomic layers. The convergence of multiple regulatory mechanisms on metabolic and translational pathways suggests a hierarchical control system where the most critical stress-responsive genes, those requiring rapid and precise expression changes, are subject to the most complex regulatory oversight.

#### 3.2.1. Integrative modelling partitions the transcriptome into five translational response programs

To integrate the multi-layer regulatory features into a coherent picture of translational control, we applied a lasso-penalised linear regression framework to predict differential STE (δSTE_M1) from the combined set of RNA features described above (**Methods**; **Figure 5A**). The number of clusters was determined by a Bayesian Information Criterion (BIC), which identified an optimal solution of five gene clusters (**Figure 5B**). Five clusters were therefore fitted using lasso regression on STE model M1, covering a total of 5,033 transcripts. The clusters differed markedly in the direction and magnitude of their δSTE distributions (**Figure 5C**), and each carried a distinct or absent functional signature as revealed by Gene Ontology enrichment analysis (**Figure 5D**).

**Figure 5.**
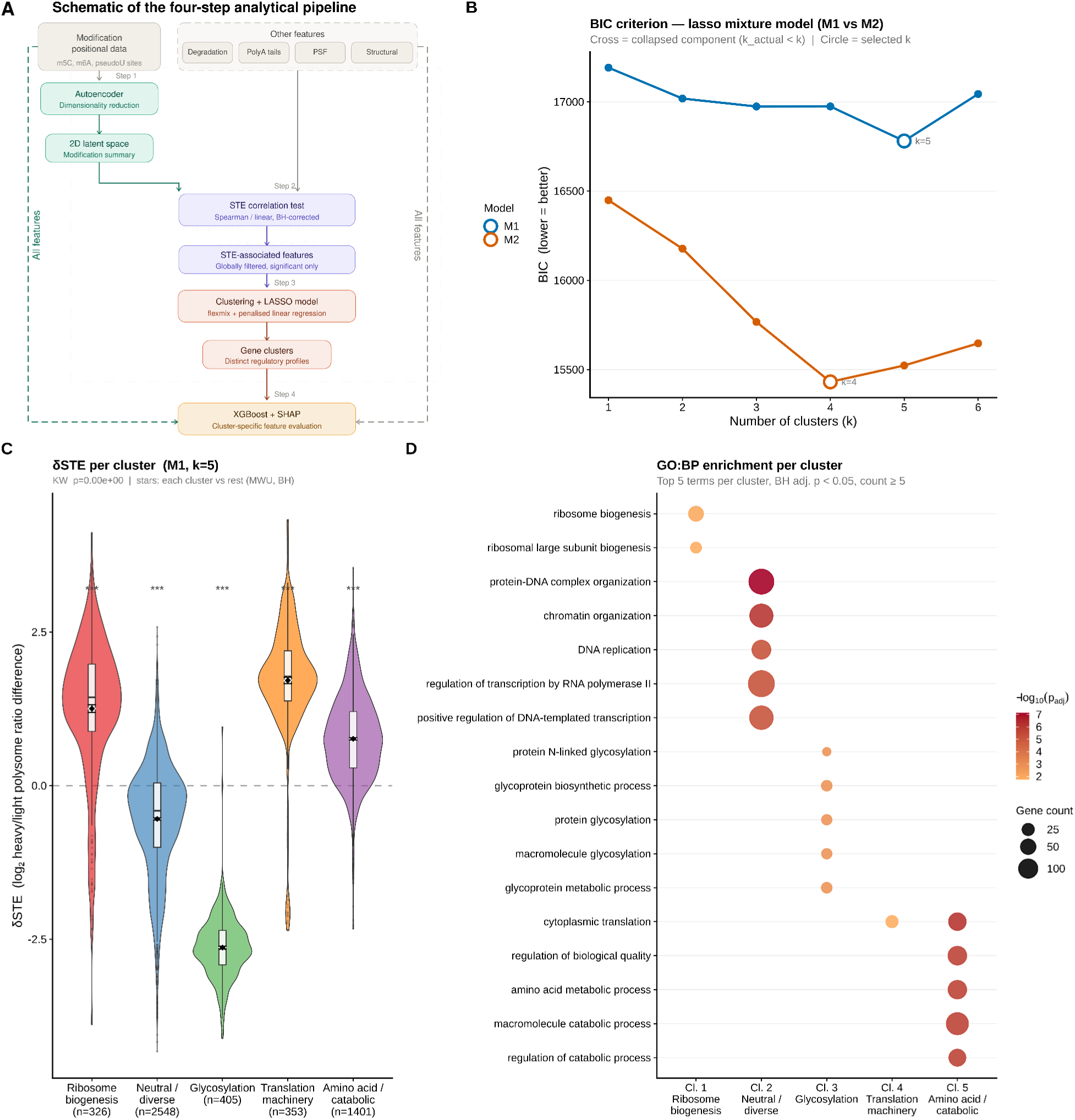
Integrative clustering identifies five translational programmes. **(A)** Schematic of the four-step analytical pipeline. Step 1: modification positional data is compressed via autoencoder to a 2D latent space. Step 2: the 2D latent space and all other features (degradation, polyA tails, PSF, structural features) enter a global STE correlation test (Spearman/linear, BH-corrected) to identify STE-associated features. Step 3: STE-associated features drive a lasso-penalised mixture of regressions (flexmix, k=5 clusters, M1). Step 4: all features re-enter an XGBoost model with SHAP for cluster-specific feature evaluation. Dashed side arrows indicate that all original features bypass directly to Step 4. **(B)** Bayesian Information Criterion (BIC) for the lasso mixture of regressions across k=1–6 clusters for Model 1 (M1, red) and Model 2 (M2, blue). Dashed vertical line marks the selected k=5 for M1. **(C)** δSTE (NS − S10) distributions across the five clusters identified by the M1 lasso mixture of regressions. Violin plots show full distributions; embedded boxplots show median and IQR; diamonds show mean ± SE. Welch ANOVA p < 10⁻¹⁰⁰. Pairwise significance brackets (BH-corrected Welch t-tests) shown above; *** p < 0.001. **(D)** GO biological process enrichment for each cluster (top 5 terms per cluster, BH adj. p < 0.05). Point size = gene count; colour = −log₁₀(adj. p). Cluster annotations at bottom indicate the dominant biological process. Cl. 1: ribosome biogenesis (n=326); Cl. 2: diverse/neutral (n=2548); Cl. 3: glycosylation (n=405); Cl. 4: translation machinery (n=353); Cl. 5: amino acid/catabolic processes (n=1401).

Three clusters showed positive δSTE. The most translationally upregulated was cluster 4 (n=353), which yielded a single, highly significant GO enrichment term: cytoplasmic translation. The selective prioritisation of the translational machinery itself as the most translationally upregulated program is striking, and suggests that an immediate response to glucose withdrawal includes the active reinforcement of translational capacity at the very onset of starvation. Cluster 1 (n=326) also showed clearly positive δSTE and was enriched specifically for ribosome biogenesis and ribosomal large subunit biogenesis, a finding that may appear counterintuitive given the high energetic cost of ribosome production, but consistent with the very early timepoint examined here (10 minutes). This effect may be a consequence of a lag before ribosome biogenesis suppression is enacted, or an active preparatory investment in translational infrastructure before longer-term transcriptional reprogramming takes over. Cluster 5 (n=1,401) showed moderately positive δSTE and was enriched for a broad set of biosynthetic and catabolic functions including cytoplasmic translation, amino acid and organic acid biosynthetic and metabolic processes, regulation of catabolic processes, and nucleocytoplasmic transport, reflecting the metabolic reprogramming toward alternative energy sources and biosynthetic programs characteristic of glucose starvation.

Cluster 3 (n=405) showed the strongest translational suppression of any cluster and was enriched for the glycosylation program, including glycoprotein biosynthesis, protein N-linked glycosylation, macromolecule glycosylation, protein modification and lipid metabolic processes. These processes are energetically costly and directly glucose-dependent (N-linked glycosylation requires dolichol-phosphate intermediates and sugar-nucleotide donors whose synthesis depends on available glucose) providing a direct metabolic rationale for their rapid and coordinated translational withdrawal upon nutrient depletion.

The largest cluster (cluster 2, n=2,548) showed near-neutral δSTE and was enriched for protein-DNA complex organisation, chromatin organisation, regulation of transcription by RNA polymerase II, DNA replication, and mitotic cell cycle progression. The absence of translational adaptation in this large cluster indicates that these housekeeping programs are not preferentially regulated at the translational level during acute starvation.

Notably, the two biological programs identified by independent layer-by-layer analysis previously (ribosomal and translational machinery on the one hand, and gluconeogenesis and metabolic reprogramming on the other) are both recovered as coherent cluster signatures in this integrative framework (**Figure 2F**), providing independent cross-validation that the multi-layer regulatory architecture described in section 2.1 reflects genuine co-regulated gene programs rather than pathway-level coincidence.

#### 3.2.2. Per-program regulatory architecture: cluster-defining features and within-cluster translational determinants

To characterise each translational program from two complementary angles, we combined a comparison of absolute feature levels across clusters, which identifies what globally distinguishes each program from the others, with an analysis of the lasso coefficients derived independently per cluster, which identifies which features predict the degree of δSTE variation among genes already assigned to that program (**Figure 6**). A cross-cluster heatmap of lasso coefficients (**Figure 6A**) reveals two cross-cutting patterns: differential model complexity, ranging from the near-single-feature architecture of cluster 3 to the multi-feature composition of cluster 4; and instances of antagonistic directionality, where the same feature predicts opposite translational outcomes in different programs (most strikingly, CDS length is the sole positive coefficient in cluster 3 yet the largest negative coefficient in cluster 4, and RDT-fraction degradation is negative in cluster 1 but positive in clusters 4 and 5). To assess whether non-linear feature interactions provide additional explanatory power, we also fitted an independent XGBoost model per cluster and derived SHAP values for feature attribution; the top SHAP features closely mirror the top lasso coefficients in all five clusters, confirming that the linear framework captures the dominant regulatory axes.

**Figure 6.**
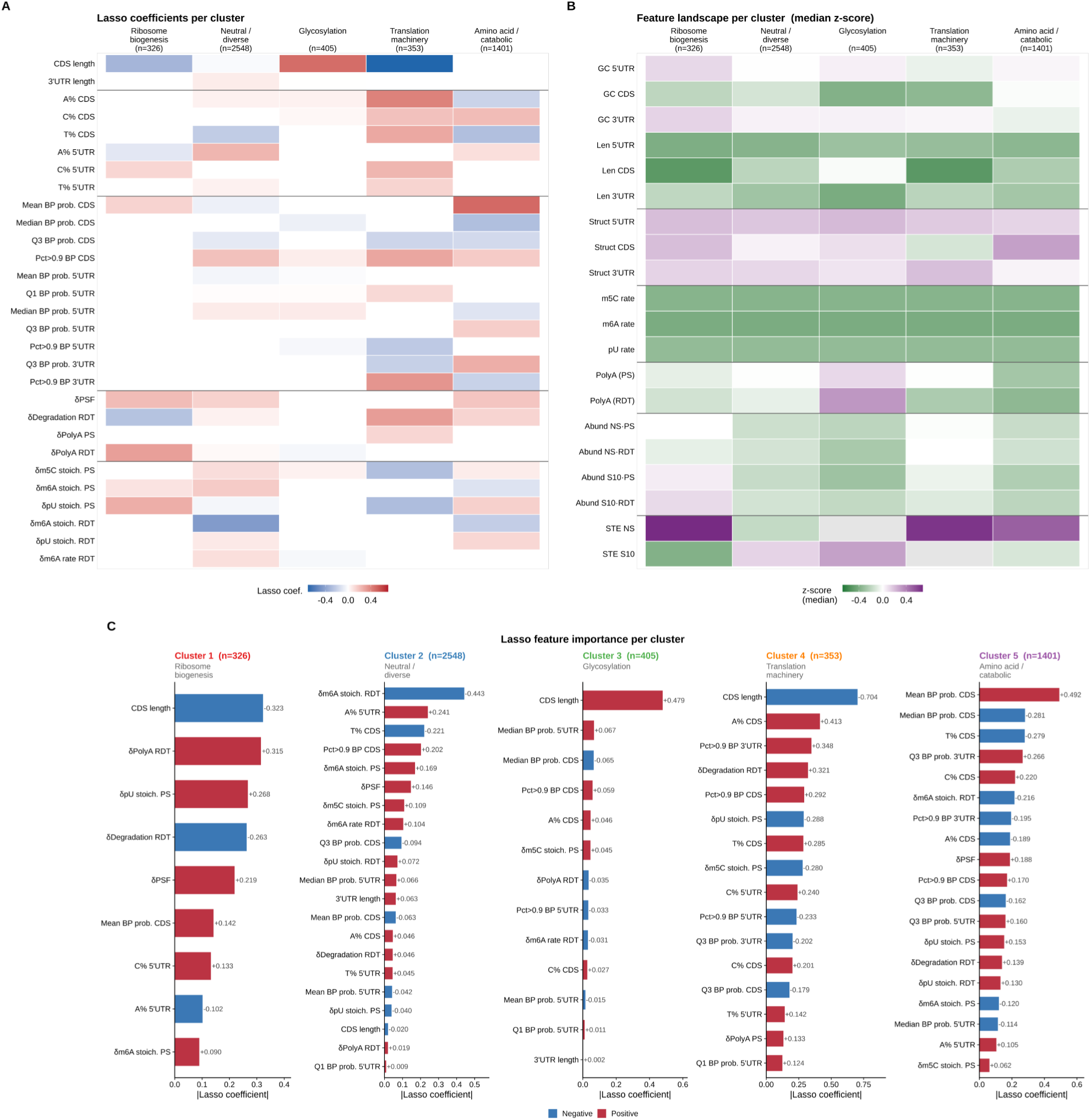
Feature architecture of the five translational programmes. (A1–A6) Per-feature violin plots for six selected features showing distributions across the five clusters. Diamonds indicate mean ± SE. Global ANOVA significance (BH-corrected, Welch ANOVA) shown above each panel; pairwise significance brackets (BH-corrected) indicate the most significant pairs. A1: CDS length (log₁₀). A2: 3’UTR length (log₁₀). A3: mean base-pairing probability (CDS). A4: δPSF. A5: δdegradation rate (RDT). A6: δpolyA tail length (RDT). **(B)** Lasso coefficient heatmap for the five-cluster M1 model. Each row is a feature; each column is a cluster. Colour encodes the lasso coefficient (red = positive association with δSTE; blue = negative). Features are grouped by category (transcript architecture, nucleotide composition, RNA secondary structure, dynamic regulatory features, RNA modification stoichiometry). Horizontal lines separate categories. Only features with at least one non-zero coefficient are shown. **(C1–C5)** Per-cluster lasso bar charts showing the non-zero lasso coefficients ranked by magnitude. Red bars indicate positive coefficients (feature values associated with higher δSTE); blue bars indicate negative coefficients. Coefficient values are annotated. Cluster biological labels and gene counts shown in the panel titles.

**Cluster 1** (ribosome biogenesis, n = 326, moderately positive δSTE) is globally distinguished not by an elevated feature but by an attenuated response across the dynamic regulatory layers. While all clusters show poly(A) tail shortening in the RDT fraction, loss of m6A, and loss of pseudouridine in the polysome fraction under starvation, these changes are far less pronounced in cluster 1 (mean Δpoly(A) RDT = −0.91 nt *versus* −2.6 to −3.3 nt across other clusters; **Figure 6B**). Cluster 1 genes also carry shorter coding sequences (mean 1,173 nt), comparable to the most translationally upregulated cluster. Interestingly however, within the program the within-cluster lasso identifies a regulatory axis centred on RNA modification and mRNA stability: pseudouridine gain in the polysome fraction and poly(A) tail maintenance in the RDT fraction are the strongest positive predictors of δSTE (coefficients +0.268 and +0.315 respectively), while increased RDT-fraction degradation is the second strongest negative predictor (−0.263) (**Figure 6C**). Together these coefficients indicate that within the ribosome biogenesis program, translational upregulation is most pronounced in genes that simultaneously gain pseudouridine in the translating pool, retain longer poly(A) tails in the ribosome-free fraction, and are stabilised rather than degraded there.

**Cluster 2** (functionally diverse, n = 2,548, near-neutral δSTE) and **Cluster 5** (amino acid metabolism and catabolic processes, n = 1,401, moderately positive δSTE) share a global feature profile distinguished by elevated δPSF (mean 0.322 and 0.333, respectively) without any strong distinguishing static architectural feature (**Figure 6B**). Their within-cluster lasso models, however, reveal distinct molecular axes. In cluster 2, the dominant within-cluster predictor is m6A stoichiometry in the RDT fraction (coefficient −0.443), which opposes a positive contribution from m6A in the polysome fraction (+0.169) (**Figure 6C**). The combination of these coefficients suggests a redistribution: transcripts that carry pre-existing m6A marks are more actively recruited into translation under starvation, appearing as a correlated depletion from the light ribosome pool and enrichment in the highly translating pool. Within the program, genes showing the strongest version of this shift are those most translationally upregulated. This pattern is consistent with m6A acting as a pre-deposited translational recruitment signal, read out selectively under conditions of global translational stress. In cluster 5, CDS secondary structure is the dominant within-cluster predictor: mean CDS base-pair probability carries the largest positive coefficient (+0.492), with the negative coefficient for median CDS structure (Q2_CDS, −0.281) (**Figure 6C**) suggesting that it is the presence of locally highly-paired regions within an otherwise variable CDS, rather than uniform secondary structure, that predicts translational upregulation. The negative contribution of RDT-fraction m6A (−0.216) echoes the cluster 2 pattern, consistent with m6A-marked transcripts in the ribosome-free pool being those most actively recruited into translation across both moderately upregulated programs.

**Cluster 3** (glycosylation, n = 405, most suppressed δSTE) is the most architecturally distinctive program at the global level, carrying the longest mean CDS (1,693 nt) and shortest mean 3ʹUTR (193 nt) of any cluster (**Figure 6B**). Within the cluster, the lasso model is dominated by CDS length (+0.479), the sole positive CDS length coefficient across all clusters: among glycosylation transcripts, those with longer coding sequences are the least suppressed. The median CDS base-pair probability (Q2_CDS) carries a negative coefficient (−0.065), indicating that transcripts with more uniformly structured coding regions are more strongly suppressed (**Figure 6C**). The model fits cluster 3 substantially better than any other (σ = 0.293 *versus* 0.61–0.93 elsewhere), reflecting the tight functional coherence of the glycosylation program and the correspondingly strong predictive signal of CDS architecture. SHAP attribution from the XGBoost model corroborates this: CDS length is the overwhelmingly dominant feature in cluster 3 (mean |SHAP| = 0.258), with the next-ranked feature an order of magnitude smaller (0.033), confirming that CDS architecture essentially accounts single-handedly for within-cluster translational variation in this program. Modification and stability features make modest contributions, with m5C gain in the polysome fraction contributing positively and poly(A) tail change in the RDT fraction negatively.

**Cluster 4** (cytoplasmic translation machinery, n = 353, most upregulated δSTE) presents a globally complementary profile to cluster 3: the shortest mean CDS (1,168 nt), the lowest mean CDS base-pair probability (0.582), and the highest increase in RDT-fraction degradation under starvation (**Figure 6B**). Within the cluster, CDS length carries the largest lasso coefficient of any feature in any cluster (−0.704): shorter transcripts within the translation-machinery program are most strongly upregulated, consistent with efficient elongation of compact, accessible coding sequences. Nucleotide composition features contribute substantially (A%, T%, and C% content in the CDS all carry positive coefficients of 0.41, 0.29, and 0.20 respectively), alongside positive contributions from 3ʹUTR structure and RDT-fraction degradation (**Figure 6C**). Notably, gains in m5C and pseudouridine in the polysome fraction carry negative coefficients (−0.280 and −0.288), indicating that within this strongly upregulated program, transcripts accruing fewer polysome-associated modifications are the most translationally activated, a pattern potentially reflecting negative feedback between modification-mediated quality control and translation efficiency for this gene class.

Across the five programs, two distinct regulatory logics emerge. In the two architecturally extreme clusters, the suppressed glycosylation program (cluster 3) and the strongly upregulated translation-machinery program (cluster 4), CDS length is the dominant within-cluster predictor, reflecting how transcript architecture shapes the degree of translational change within a globally defined program. In the three remaining clusters, dynamic epitranscriptomic marks and mRNA stability features drive within-cluster variation, with m6A compartmentalisation (polysome *versus* ribosome-free fraction) emerging as a recurring axis. The between-cluster comparisons establish that the translational direction of each program is set by its global feature context, while the lasso coefficients reveal the molecular levers that fine-tune individual gene responses within each. The XGBoost SHAP analysis provides independent support for this framework: the top SHAP-ranked features per cluster closely mirror the top lasso coefficients, and the between-cluster differences in model complexity observed in the lasso heatmap are recapitulated in the SHAP bar charts: CDS length dominates clusters 3 and 4 in both frameworks, while clusters 1, 2, and 5 show broader feature distributions. Finally, the two representative transcripts highlighted in section 3.1.5, YLL039C (ubiquitin) and YKR097W (PCK1), both fall within cluster 2. Their placement in the near-neutral program, despite displaying coordinated multi-layer regulatory changes at the individual level, illustrates that the combined effect of orchestrated modifications across layers can be balanced rather than amplified at the translational output level.

#### 3.2.3. CDS codon usage discriminates translational programs, and is independently confirmed by direct codon optimality indices

To determine whether the sequence composition of coding and non-coding regions contributes to the identity of the five translational programs, we computed k-mer Shannon entropy and direct nucleotide composition across the 5ʹUTR, 3ʹUTR, and CDS for all 5,033 transcripts. Between-cluster differences were assessed with ANCOVA controlling for log-transformed region length throughout.

For both UTRs, the signal was present but modest. Five-prime UTR entropy showed no significant between-cluster differences after length correction (ANCOVA, cluster term p > 0.10 at all k, BH-corrected), and direct nucleotide composition analysis revealed small but significant differences in A% and T% (ANCOVA p < 0.01, partial ηZ < 0.005), with cluster 4 having modestly higher T content and cluster 2 higher A content in 5ʹUTRs. For 3ʹUTRs, cluster 5 (amino acid and catabolic processes) had the highest AU content (median A+T = 70.9%) and cluster 3 (glycosylation) the lowest T content among the five programs (ANCOVA p < 0.01, partial η^2^ = 0.003), consistent with the known role of uridine-rich 3ʹUTR elements in recruiting decay factors in yeast (58,59). Effect sizes across all UTR composition metrics were small, however, and none of these features constitute primary determinants of program identity.

The CDS analysis yielded a qualitatively different and more interpretable result. CDS k-mer entropy was significantly higher in cluster 3 and lower in clusters 1 and 4 at every k value tested from k=1 to k=6, with the separation between programs increasing with k (ANCOVA cluster term p < 10^−7^ for all k after BH correction; partial η^2^ = 0.007-0.019; **Figure 7A**). This pattern cannot be explained by nucleotide composition alone and implicates codon-level sequence structure.

**Figure 7.**
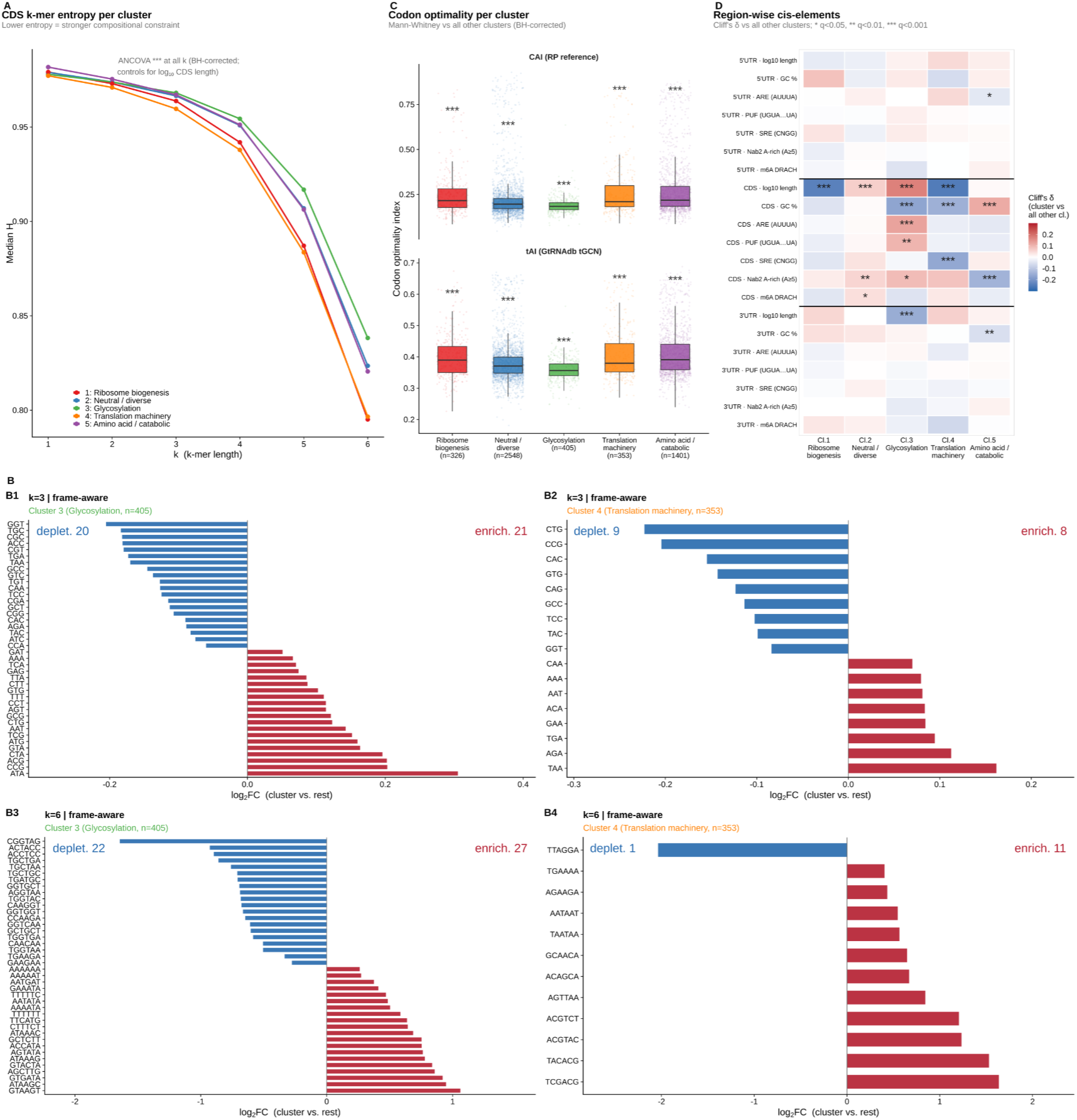
CDS sequence composition underlies cluster-specific translational programmes. **(A)** Median theoretical maximum entropy (H_theo) of CDS k-mer distributions as a function of k-mer order (k=1–6) for each cluster. Lower H_theo at high k indicates stronger compositional constraint (non-random k-mer usage). Significance of cluster differences beyond CDS length (ANCOVA, BH-corrected) is shown above each k. *** p < 0.001. Cluster 4 (translation machinery, orange) and Cluster 1 (ribosome biogenesis, red) show the strongest compositional constraint at k=5,6, while Cluster 3 (glycosylationà shows the weakest. **(B1–B4)** Frame-aware CDS k-mer enrichment (cluster vs. all other genes, Fisher exact test, BH-corrected) for clusters 3 and 4. Only k-mers starting at codon positions (1, 4, 7, …) are counted. B1: k=3 (codon-level), Cluster 3 (Glycosylation). B2: k=3, Cluster 4 (Translation machinery). B3: k=6 (codon-pair level), Cluster 3. B4: k=6, Cluster 4. Bars show log₂ fold-change; red = enriched in cluster, blue = depleted. Numbers of significant enriched/depleted k-mers are annotated. Top 15 enriched and depleted k-mers shown per panel. **(C)** Codon Adaptation Index (CAI, top) and tRNA Adaptation Index (tAI, bottom) per cluster (jitter + boxplot; n = 326–2,548 genes per cluster). Cluster 3 (glycosylation) shows the lowest codon optimality of any programme by both measures; clusters 4 and 5 carry the most optimised coding sequences. Stars indicate Mann-Whitney tests of each cluster against all other clusters combined (BH-corrected); *** p < 0.001 for all ten cluster × metric comparisons. **(D)** Region-wise scan for cis-regulatory sequence composition and motifs (GC content, AU-rich elements [ARE], Puf-binding motifs [PUF], Smaug/Vts1 response elements [SRE], Nab2 A-rich tracts, and m6A DRACH consensus sites) across the 5′UTR, CDS, and 3′UTR of each cluster, expressed as Cliff’s δ relative to all other clusters combined (Mann-Whitney, BH-corrected). Of 105 cluster × region × feature tests, the significant signals are concentrated almost entirely in the CDS row block with only sporadic, small-effect UTR hits (5′UTR GC in cluster 1, 3′UTR length in clusters 2 and 3), confirming that the compositional divergence between programmes is a CDS-specific phenomenon rather than a UTR-driven one. * q < 0.05, ** q < 0.01, *** q < 0.001.

To test this interpretation, we performed a frame-aware k-mer enrichment analysis, restricting the comparison to k-mers starting at codon positions (positions 1, 4, 7,… in each CDS), which isolates true codon usage from non-frame-specific sequence context. At k=3, this analysis directly identifies differentially used codons between each cluster and the rest (Fisher’s exact test, BH correction; **Figure 7B1,B2**). Cluster 4 (translation machinery) was enriched for A-rich codons: AAA and GAA [Lys and Glu, the two most abundant amino acids in yeast ribosomal proteins(60), AGA (Arg), and CAA (Gln)], together with the TAA stop codon, which is the preferred stop codon of highly expressed genes in yeast(61,62). Cluster 3 (glycosylation) showed the opposite profile: enrichment of non-optimal codons including ATA (Ile), CTA (Leu), ACG (Thr), and GTA (Val), which are among the rarest codons in the yeast high-expression gene set(51).

Extending the frame-aware analysis to k=6, which counts consecutive codon pairs and tests di-codon usage, recapitulated and extended these findings (**Figure 7B3,B4**). Cluster 4 was enriched for repeated A-rich codon pairs (AGAAGA, Arg-Arg; AATAAT, Asn-Asn), consistent with the repetitive amino acid stretches characteristic of ribosomal proteins. Cluster 3 was enriched for TTTTTT (Phe-Phe di-codon) and depleted for GGTGGT (Gly-Gly) and CAACAA (Gln-Gln) pairs found preferentially in well-adapted coding sequences.

The two indices give concordant genome-wide rankings (CAI-tAI Spearman ρ = 0.93) and both correlate positively with δSTE across the full transcriptome (CAI ρ = 0.31, p < 10^−15^; tAI ρ = 0.34, p < 10^−15^), indicating that transcripts with more codon-optimised coding sequences are precisely those whose translation is most sustained under starvation. At the cluster level, cluster 3 (glycosylation) has the lowest codon optimality of any program by both measures relative to all other clusters combined (CAI: Cliff’s δ = −0.29, q = 1 × 10^−21^; tAI: δ = −0.33, q = 5 × 10^−27^), while clusters 4 (translation machinery) and 5 (amino acid/catabolic) carry the most optimised coding sequences (CAI: δ = +0.13 and +0.24 respectively, both q < 10^−4^; **Figure 7C**).

To quantify codon optimality directly, independently of the entropy framework, we computed two independent codon optimality indices for each transcript: the Codon Adaptation Index [CAI;(51)], using the 130 cytoplasmic ribosomal-protein genes as the yeast reference set, and the tRNA Adaptation Index [tAI;(52)], derived from the GtRNAdb sacCer3 tRNA gene-copy numbers(63). The two indices give concordant genome-wide rankings (CAI-tAI Spearman ρ = 0.93) and both correlate positively with δSTE across the full transcriptome (CAI ρ = 0.31, p < 10^−15^; tAI ρ = 0.34, p < 10^−15^), indicating that transcripts with more codon-optimised coding sequences are precisely those whose translation is most sustained under starvation. At the cluster level, cluster 3 (glycosylation) has the lowest codon optimality of any program by both measures relative to all other clusters combined (CAI: Cliff’s δ = −0.29, q = 1 × 10^−21^; tAI: δ = −0.33, q = 5 × 10^−27^), while clusters 4 (translation machinery) and 5 (amino acid/catabolic) carry the most optimised coding sequences (CAI: δ = +0.13 and +0.24 respectively, both q < 10^−4^; **Figure 7C**). The agreement between CAI and tAI confirms that the codon usage divergence between programs is not an artefact of the entropy metric or the choice of reference set, but reflects a genuine, two-reference-validated difference in how well each cluster’s transcripts are matched to the yeast tRNA pool.

Finally, a complementary region-wise scan for known *cis*-regulatory sequence elements, adenosine-uridine-rich elements (AREs), Puf-family binding motifs, stress-response elements (SREs), and m6A DRACH motifs, similarly found no significant cluster-specific enrichment in either UTR across all five clusters (**Figure 7D**). The significant signals were confined to the CDS, where log_10_(length) and GC content showed cluster-specific differences consistent with the k-mer entropy analysis above.

#### 3.2.4. Convergent *cis*-regulatory features define the translational fate of the glycosylation program

The codon optimality contrast between cluster 3 and the other programs raises the question of whether additional *cis*-regulatory features converge on the same axis. We therefore systematically scored and tested each cluster’s transcripts against a matched background for four further feature families: initiation context, upstream open reading frames, start-codon RNA structure, and experimental RNA-binding protein (RBP) regulon membership (see **Methods**).

**Initiation context.** Extended Kozak context was scored as a position-weight match over the −6 to +4 window around the initiator AUG. Cluster 3 carries the weakest Kozak context of any cluster (Cliff’s δ = −0.217 *vs.* background, q = 8×10^−12^), with the deficit concentrated at the functionally most critical positions (−3 and +4(53); **Figure 8A1**). Cluster 4 and cluster 5 carry the strongest contexts (δ = +0.101,q = 2.7×10^−3^ and δ =+0.089, q = 2.7×10^−6^ respectively). To test whether initiation context tunes the starvation response within each program, we computed the partial Spearman correlation between Kozak strength and δSTE controlling for transcript length. This coupling is present and significant in cluster 5 (ρ = 0.20, q = 2 × 10^−13^) and in the diverse background cluster 2 (ρ = 0.10, q = 7 × 10^−7^), but is absent in cluster 3 (ρ = −0.03, n.s.; **Figure 8A2**): in the glycosylation program, ground-state initiation efficiency no longer predicts the magnitude of translational repression, even though this cluster carries the weakest contexts on average. This decoupling suggests that the translational response of cluster 3 is imposed by a downstream layer rather than set by initiation.

**Figure 8.**
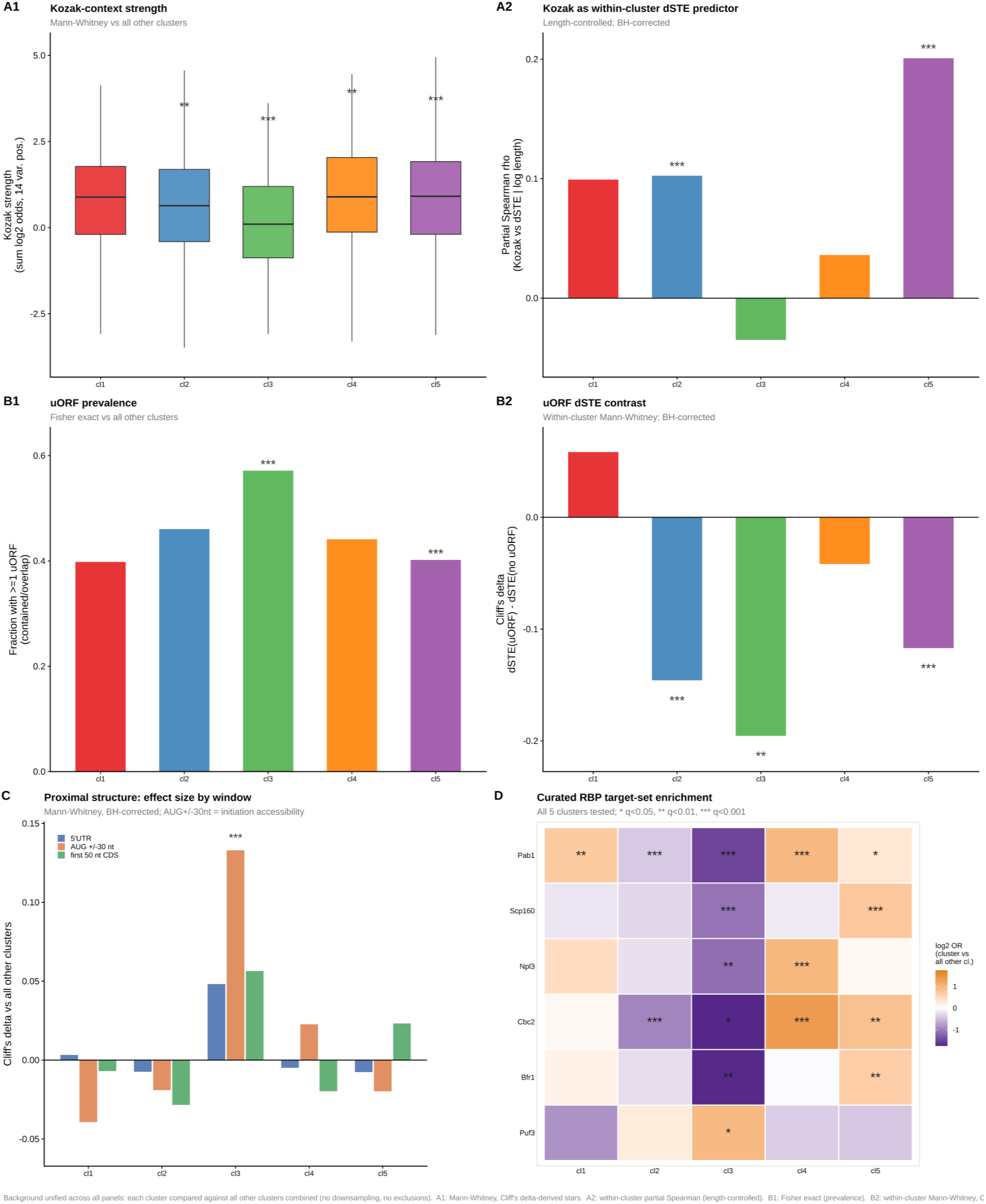
Convergent cis-regulatory features distinguish the glycosylation programme from the rest of the translational landscape. (A1) Kozak-context strength (summed log₂ odds over the 14 variable positions of the −10 to +7 window around the initiator AUG) per cluster (boxplots). Cluster 3 (glycosylation) carries the weakest Kozak context of any programme (Cliff’s δ = −0.22 vs. all other clusters, q = 8.1 × 10⁻¹Z); clusters 4 and 5 carry the strongest (δ = +0.10, q = 2.7 × 10⁻3 and δ = +0.09, q = 2.7 × 10⁻⁶, respectively). ** q < 0.01, *** q < 0.001. **(A2)** Kozak strength as a within-cluster predictor of δSTE (length-controlled partial Spearman ρ, BH-corrected across clusters). This coupling is strongest in cluster 5 (ρ = 0.20, q = 2.3 × 10⁻¹3) and present in the diverse background cluster 2 (ρ = 0.10, q = 6.8 × 10⁻⁷), but is absent in cluster 3 (ρ = −0.03, n.s.): in the glycosylation programme, initiation-context strength no longer predicts the magnitude of translational repression, despite carrying the weakest contexts on average. **(B1)** Fraction of transcripts carrying at least one upstream open reading frame (uORF; contained or main-AUG-overlapping) per cluster (Fisher exact vs. all other clusters, BH-corrected). Cluster 3 is the most uORF-laden programme (57.1% vs. 43.7% background, q = 1.7 × 10⁻⁶); cluster 5 is depleted (40.2% vs. 46.5%, q = 1.6 × 10⁻⁴). *** q < 0.001. **(B2)** Within-cluster δSTE contrast between uORF-bearing and uORF-free transcripts (Cliff’s δ, Mann-Whitney, BH-corrected). Negative values indicate translational repression by uORFs. The repression signal is strongest in cluster 3 (δ = −0.20, q = 1.5 × 10⁻3) and cluster 2 (δ = −0.15, q < 10⁻⁸), and is absent in the high-δSTE translation-associated clusters 1 and 4 (both n.s.), distinguishing this population-level pattern from the GCN4-style translational de-repression seen under canonical amino acid starvation. **(C)** Proximal RNA secondary structure (mean base-pairing probability from RNAfold predictions) summarised as Cliff’s δ relative to all other clusters across three windows per transcript: 5′UTR, AUG ± 30 nt (initiation accessibility), and the first 50 nt of the CDS. Of fifteen region × cluster tests, only cluster 3 reaches significance, and only in the AUG ± 30 nt window (δ = +0.13, q = 4.9 × 10⁻⁴), indicating excess structure specifically at the initiation site. *** q < 0.001. **(D)** Curated experimental RNA-binding protein (RBP) target-set enrichment for a focused panel of six RBPs (Pab1, Scp160, Npl3, Cbc2, Bfr1, Puf3; from the Hogan et al. 2008 RIP-chip atlas), expressed as log₂ odds ratio of cluster membership relative to all other clusters (Fisher exact, BH-corrected across all 5 clusters × 42 RBPs tested). Cluster 3 is significantly depleted for the targets of the major stabilising and export-associated RBPs (Pab1, Scp160, Npl3, Cbc2, Bfr1), while showing nominal enrichment for the destabilising Puf3 regulon. Clusters 4 and 5 show the mirror enrichment pattern for the same poly(A)/export-associated sets. * q < 0.05, ** q < 0.01, *** q < 0.001.

**Upstream open reading frames.** uORFs were called as AUG-to-stop reading frames in each annotated 5ʹUTR. Cluster 3 is the most uORF-laden program, both in prevalence (enriched relative to background; −log_2_ odds ratio = 0.81, q < 10^−3^) and in uAUG density (q < 10^−3^; **Figure 8B1**). Clusters 1 and 5 are depleted for uORFs. Crucially, at the transcriptome level uORF-bearing transcripts show consistently lower δSTE (median −0.12 *versus* +0.16 for uORF-free; Cliff’s δ = −0.15; uAUG density ρ = −0.12), and this repression signal is strongest in the most uORF-dense cluster (cluster 3: δ = −0.2, q = 1.5 × 10^−3^) and absent in the high-δSTE translation clusters (clusters 1 and 4, both n.s.; **Figure 8B2**). Notably, this is a population-level uORF repression pattern, unlike the well-characterised GCN4 mRNA, whose uORFs permit translational induction under amino acid starvation(7): uORFs systematically reduce, rather than boost, translation under starvation across the bulk transcriptome, with cluster 3 as the extreme case.

**Start-codon RNA structure.** Base-pair probability was computed from RNAfold predictions sliced into three windows per transcript (5ʹUTR, AUG ± 30 nt, early CDS). Of fifteen region × cluster tests, the only significant result is cluster 3, whose start-codon window is more base-paired than background (Cliff’s δ = +0.13, q = 5 × 10^−4^; **Figure 8C**). A metagene profile of P(paired) over offsets −100 to +150 around the AUG localises the excess pairing specifically over the initiation site, with 5ʹUTR and CDS windows flat. No other cluster shows a significant start-codon structure effect; cluster 5 trends in the opposite direction. Together with the uORF data, the start-codon structure finding identifies the initiation step, and not elongation, as the primary site of translational control in cluster 3.

**Experimental RBP regulon membership.** To test whether the *cis*-feature convergence is mirrored by experimental evidence for RBP regulation, we tested cluster membership against 42 curated RBP regulon sets derived from the Hogan *et al.* (2008)(54) RIP-chip atlas (42 RBPs with at least one target at 1% SAM-FDR; Methods), with independent cross-validation using the Gerber *et al.* (2004) (55) Puf1-5 sets. Of 168 cluster × RBP Fisher exact tests (BH-corrected), 18 are significant at q < 0.05 (**Figure 8D**). Cluster 3 is significantly depleted for the targets of the major poly(A)-binding, stabilising, and nuclear-export RBPs, such as Pab1 (log_2_ OR = −1.87, q = 1.7 × 10^−13^), Npl3, Scp160, Puf4, Bfr1, and Pub1, while being enriched for the Puf3 regulon (log_2_ OR = +1.25), whose targets are preferentially destabilised(54). Clusters 4 and 5 show the mirror enrichment for the same poly(A)/export sets. The experimental RBP target layer is orthogonal to the motif scan (which uses sequence patterns rather than *in vivo* binding) and to the codon-optimality and structure analyses, yet it recovers the same cluster 3 repressor signature from a completely independent source.

**Convergence across feature families.** Assembling the effect sizes from all five feature families into a single oriented matrix (**Figure 9A**) makes the convergence explicit. Cluster 3ʹs column is uniformly negative across every axis: lowest codon optimality, weakest Kozak context, highest uORF load, most structured start codon, and most depleted of stabilising RBP targets, and its δSTE is uORF-coupled yet Kozak-decoupled. Cluster 4 is the consistent positive mirror on the codon and RBP axes. That four feature families estimated by entirely independent methods all point to the same cluster argues that the glycosylation program’s translational repression is not driven by a single regulatory element but is written into multiple independent layers of the *cis*-regulatory architecture. The convergence of four independent feature families on the same functional cluster, and the failure of any single feature to account fully for the observed suppression, suggests that the glycosylation program’s translational fate is the result of deep, multi-layered *cis*-regulatory encoding rather than the action of a single dominant mechanism, a point we return to in the **Discussion**.

**Figure 9.**
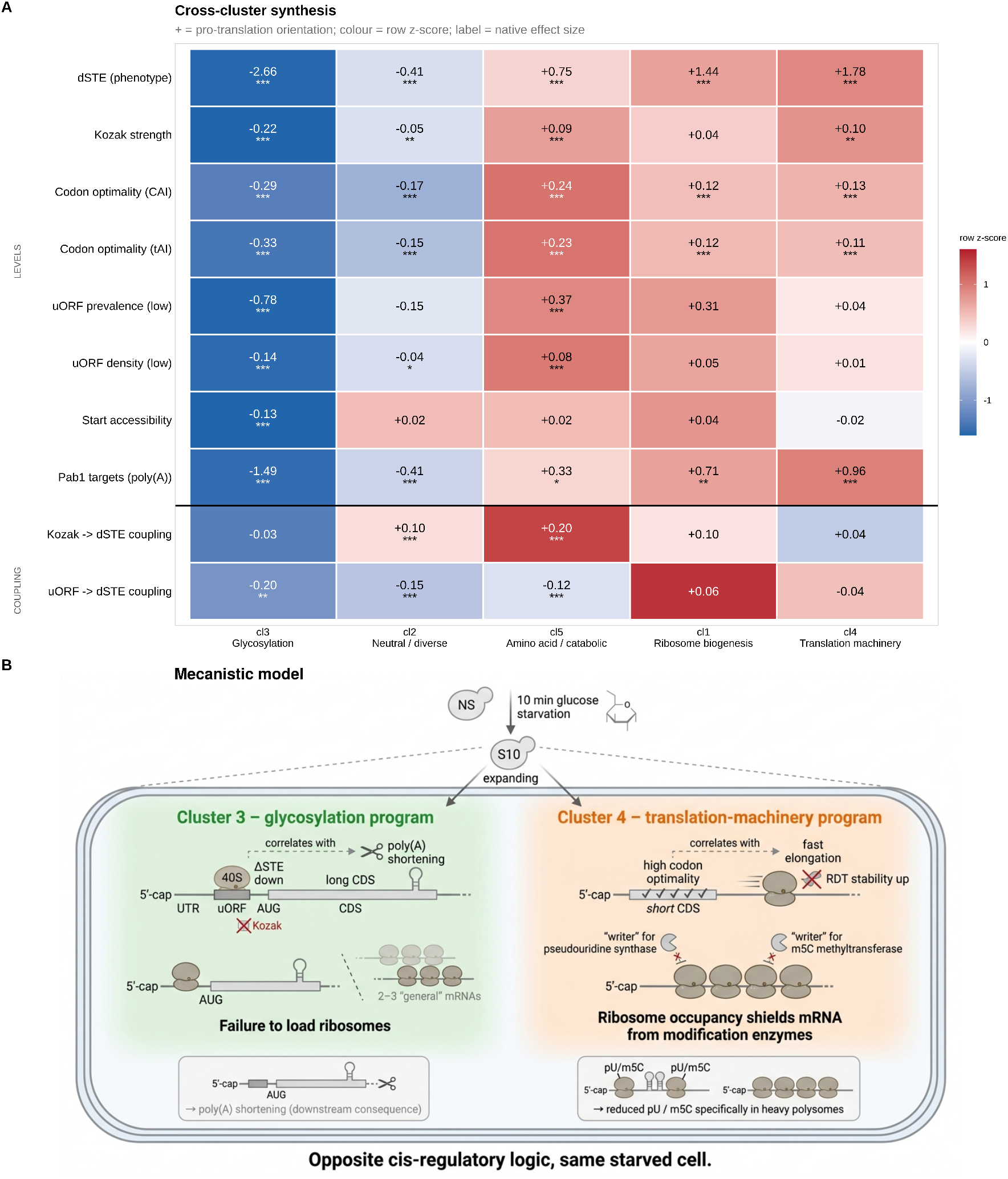
Cross-cluster synthesis of feature-phenotype associations and a mechanistic model contrasting the glycosylation and translation-machinery programs. **(A)** Heatmap summarising, for each of the five clusters, the association between regulatory features and the starvation-induced change in translation efficiency (dSTE). Clusters are ordered left to right by increasing dSTE, from most suppressed (cl3, glycosylation) to most upregulated (cl4, translation machinery), with the near-neutral (cl2), amino-acid/catabolic (cl5), and ribosome-biogenesis (cl1) clusters in between. Cell colour encodes the row z-score (i.e. each feature’s association strength scaled across the five clusters, to allow comparison of relative direction and magnitude across a row); the printed value is the native, unscaled effect size for that feature-cluster pair, and asterisks denote statistical significance (*//*; see Methods for exact thresholds). The upper block (LEVELS) reports each cluster’s characteristic feature values — dSTE itself, Kozak strength, codon optimality (CAI, tAI), uORF prevalence/density, start-codon accessibility, and Pab1 (poly(A)-binding protein) target status — relative to the other clusters. The lower block (COUPLING, below the horizontal divider) reports how strongly Kozak strength and uORF density are associated with dSTE within each cluster, i.e. how much each feature actually predicts translational response in that cluster, as distinct from the feature’s typical level shown above. **(B)** Proposed mechanistic model contrasting Cluster 3 (glycosylation program) and Cluster 4 (translation-machinery program), shown within a single yeast cell 10 minutes into glucose starvation (S10), starting from the non-starved (NS) state. Both programs co-occur within the same starved cell and represent opposite cis-regulatory logic. In Cluster 3 (green), transcripts combine a long CDS with an upstream ORF (uORF) preceding the main AUG; despite a favourable Kozak context, the 40S scanning subunit fails to efficiently reach the main start codon, so these transcripts fail to load ribosomes relative to the general translating pool. This translational exclusion is the primary outcome; progressive poly(A) shortening is shown as a downstream consequence rather than the driving event. In Cluster 4 (orange), transcripts combine a short CDS with high codon optimality, supporting fast, smooth ribosome elongation and increased RDT-fraction (light-polysome) stability. The resulting dense ribosome occupancy is proposed to sterically shield these transcripts from pseudouridine-synthase and m5C-methyltransferase “writer” enzymes, so that pseudouridine and m5C marks are specifically reduced on heavily ribosome-loaded (PS-fraction) transcripts. Together, the two programs illustrate how opposite cis-encoded architectures — an initiation-limiting versus an elongation-favouring configuration — drive divergent post-transcriptional fates within the same acutely starved cell.

## 4. Discussion

### 4.1. Direct RNA Sequencing provides a unique readout of multi-omic RNA regulatory layers

A central practical advantage of the direct RNA sequencing approach used here is that it allowed transcript abundance, mRNA degradation, poly(A) tail length, and the stoichiometry of three distinct RNA modifications (m5C, m6A, pseudouridine) to be measured simultaneously, from the same molecules, in both the light and heavy polysome-associated mRNA fractions. Each of these regulatory layers has traditionally required its own dedicated experimental protocol, RNA-seq for abundance, metabolic labelling or actinomycin-D chase for degradation, TAIL-seq or PAL-seq for poly(A) length, and antibody- or chemistry-specific protocols for each individual modification, making it difficult to relate them to one another within the same biological sample, let alone the same molecule. By contrast, direct nanopore sequencing of native RNA preserves all of this information in a single long read(64,65), removing both the technical noise introduced by combining independently generated datasets and the biological noise introduced by comparing measurements taken in different cells, at different times, or under subtly different growth conditions. This is the foundation that made the multi-omic integration reported here possible.

### 4.2. Individual layers hint at regulatory structure but do not, on their own, resolve translational programs

When examined one layer at a time (**Results, Section 3.1**), several of these regulatory axes showed clear and statistically robust changes under glucose starvation, and in some cases these changes were concentrated in recognisable functional pathways: transcripts encoding ribosomal proteins and translation factors were more stable in non-starved cells, while gluconeogenic and stress-protective transcripts showed the opposite pattern; poly(A) tail dynamics and modification stoichiometry shifted in a fraction- and pathway-dependent manner (**Figure 1**). However, the relationship between any single layer and translational efficiency itself was inconsistent and often counterintuitive: transcripts stabilised under starvation were not necessarily more efficiently translated, and transcripts with the largest changes in modification stoichiometry did not map cleanly onto a single direction of translational consequence. This layer-by-layer analysis shows that abundance, stability, polyadenylation, and modification state are not independent, redundant readouts of the same regulatory event, and that no single layer, nor an informal combination of several, offered a coherent account of how translational efficiency itself was being set. The uni-omic results pointed toward coordinated regulation without specifying its structure.

### 4.3. RNA multi-omic integration resolves five distinct translational programs

This structure became apparent only when all measured features were combined within a single lasso-penalised mixture-of-regressions model predicting δSTE (**Results, Section 3.2**). This joint model partitioned the 5,033 analysed transcripts into five translational programs with sharply distinct δSTE distributions and well-defined functional identities (**Figure 4C-D**): a translation-machinery program strongly upregulated under starvation, a ribosome-biogenesis program more modestly upregulated, a glycosylation program strongly repressed, a moderately upregulated amino-acid/catabolic program, and a large, near-neutral housekeeping program. While the translation-machinery and glycosylation program were already visible in the layer-by-layer analysis, it could not provide a quantitative, transcriptome-wide partition: it could not assign individual transcripts to programs with defined boundaries, could not resolve how many distinct regulatory programs existed, could not separate the near-neutral housekeeping bulk of the transcriptome from genuinely regulated transcripts, and established only a qualitative association between each layer and pathway identity rather than a quantitative relationship to translational efficiency itself.

The joint lasso mixture model resolved all four of these: it recovered the same translation-machinery/glycosylation contrast as the most extreme pair of programs (independently re-deriving, rather than simply assuming, the two pathways already implicated by the uni-omic analysis), but additionally resolved three further programs (ribosome biogenesis, amino-acid/catabolic, and housekeeping) that were not cleanly separable from one another or from the two extremes using any single layer, assigned every transcript a specific cluster membership, and directly quantified each program’s relationship to δSTE. This is the first piece of evidence that integration provides more than the sum of its parts: not the discovery of a contrast invisible to uni-omic analysis, but the resolution of that contrast into a specific, quantitative, transcript-resolved structure that no single layer could deliver on its own.

### 4.4. RNA multi-omic modelling framework

Our modeling strategy was designed to specifically capture the molecular response to glycemic stress rather than constitutive differences between conditions. Instead of modeling starved and non-starved measurements separately, we used feature-wise differences across all omics layers. This transformation focuses the analysis on stress-induced changes and reduces the influence of baseline variability between genes. However, using feature-wise differences inevitably removes information contained in the absolute measurements under each condition. Consequently, regulatory mechanisms that depend on baseline molecular states, rather than on the magnitude of the stress response, may not be detected. Our post-clustering analysis of each of the features therefore further characterized the clusters.

To account for the heterogeneity of translational regulation, we used a non-supervised clustering strategy based on finite mixture regression models, in which cluster assignment is optimized according to the relationship between δSTE and the multi-omic predictors rather than the similarity of the predictors themselves. Consequently, the identified clusters should be interpreted as predictive regulatory programs, representing groups of genes that share similar associations between molecular features and translational response. They do not necessarily correspond to intrinsic biological groupings that would emerge from an unsupervised clustering of the multi-omic data. Different clustering strategies or alternative response variables could therefore produce different partitions of the transcriptome.

Finally, XGBoost models were trained independently within each cluster, and SHAP values were used to quantify feature contributions. This two-step approach captures nonlinear relationships and interactions that are not modeled by the initial linear mixture regression, while SHAP improves model interpretability. However, SHAP feature importance reflects predictive contribution rather than causality and may be affected by correlations among omics features. Consequently, the identified regulators should be interpreted as candidate explanatory variables that warrant experimental validation rather than definitive mechanistic drivers.

### 4.5. Same features can have opposite regulatory consequences in different programs

The cluster-specific coefficients estimated by the joint model (**Figure 5A**) make this non-additivity concrete and quantifiable. Several features carry opposite-signed coefficients across clusters, meaning the same molecular measurement predicts translational upregulation in one program and repression in another. The clearest example is CDS length, the single largest-magnitude predictor in the entire model: longer coding sequences predict *reduced* repression in the glycosylation program (cluster 3, β = +0.48) but predict the *strongest* translational *upregulation* of any feature in the translation-machinery program (cluster 4, β = −0.70, the largest-magnitude coefficient anywhere in the model). A transcriptome-wide regression of δSTE on CDS length that ignored cluster membership would partially cancel these opposing contributions, understating the role CDS length plays in either program considered on its own.

This sign reversal is not confined to single features in isolation; it extends to pairs of features whose *joint* regulatory relationship flips across clusters. CDS length and RDT-fraction degradation rate change illustrate this directly: in cluster 1 (ribosome biogenesis), both features carry negative coefficients (CDS length β = −0.32, RDT degradation β = −0.26): shorter, less-stabilised transcripts are associated with greater translational upregulation, a single coherent direction of effect. In cluster 4 (translation machinery), the two features decouple: CDS length remains strongly negative (β = −0.70), but RDT degradation flips to strongly positive (β = +0.32): within this program, transcripts whose RDT-fraction stability *increases* under starvation are now the ones most translationally upregulated, the opposite relationship to cluster 1. A model that treated CDS length and RDT degradation as having a fixed, additive relationship to translational efficiency would necessarily get one of these two programs wrong, because the pair’s *joint* regulatory sign is not a fixed property of the two features but a property of the cluster they are being evaluated within.

A complementary pattern further illustrates what is gained by integration: features with weak, easily-overlooked marginal effects can become strong, well-supported predictors once cluster structure is incorporated. RDT-fraction m6A stoichiometry change is the clearest case. Across the full transcriptome, ignoring cluster membership entirely, its marginal correlation with δSTE is modest (Spearman ρ = −0.13, p = 1.9 × 10^−21^), detectable given the large sample size, but not the kind of effect size that would mark a feature as a central regulatory determinant. Within cluster 2 alone, however, the same correlation more than doubles in magnitude (ρ = −0.29, p = 1.3 × 10 ^−50^), and the lasso model independently assigns RDT m6A change the single largest-magnitude coefficient of any feature in that cluster (β = −0.44) while shrinking it to exactly zero in three of the other four clusters. A feature-by-feature screen applied to the full transcriptome would have ranked RDT m6A change well below CDS length, degradation rate, or codon-related features in apparent importance; only by allowing its effect to be cluster-specific does its true, substantially larger role within a defined transcript population become visible.

Together, these observations show that the regulatory architecture uncovered here is not additive: the relationship between a given molecular feature and translational efficiency depends on the broader combination of features and the cluster membership they jointly imply, sometimes reversing in sign and sometimes appearing negligible until restricted to the right transcript population. This is the central methodological argument of the present work: that resolving the post-transcriptional regulatory logic of the starvation response required modelling these layers jointly, and that any approach examining transcript abundance, stability, polyadenylation, or modification state independently, however comprehensive each individual measurement, would have systematically understated or entirely missed this structure.

### 4.6. The *cis*-regulatory architecture suggests program-specific mechanistic models

The *cis*-regulatory features described in **Results Section 3.2** characterise the sequence and trans-regulatory architecture of each cluster’s transcripts independently of how δSTE is predicted within each program. Asking how these *cis*-features relate to the dynamic features the lasso model does use as predictors reveals coherent, program-specific mechanistic hypotheses that connect pre-encoded sequence architecture to the post-transcriptional dynamics observed under starvation. We present these connections as a proposed mechanistic model rather than an established causal chain, as the evidence is correlational throughout (**Figure 8B**).

Within the glycosylation program (cluster 3), the lasso model predicts the change in translation efficiency primarily from CDS length, CDS base-pairing probability, and poly(A) shortening in the light-ribosome fraction. Within this cluster, uORF density correlates negatively with both CDS length (ρ = −0.17) and poly(A) shortening (ρ = −0.12). This suggests that the *cis*-encoded burden of upstream open reading frames impedes productive ribosome scanning to the main AUG, reducing polysome loading; transcripts that fail to load ribosomes efficiently accumulate in the light polysome fraction, where extended exposure to cytoplasmic deadenylases drives disproportionate poly(A) loss. In this model, the cis architecture sets initiation efficiency, which jointly determines both δSTE and the extent of poly(A) loss downstream. Consistent with this, Kozak strength does not predict δSTE within cluster 3 (partial ρ = −0.03, n.s.; **Figure 7A2**): even transcripts with stronger-than-average initiation contexts are not rescued from repression, indicating that uORF burden and start-codon structure constitute the dominant, Kozak-overriding barriers in this program.

Within the translation-machinery program (cluster 4), the lasso predicts the change in translation efficiency from short CDS length, increased RDT-fraction stability, and reduced polysome-fraction pseudouridine and m5C. Here codon optimality (CAI) correlates positively with RDT-fraction stability (ρ = +0.16) and with both modification changes within the cluster (pU: ρ = +0.18; m5C: ρ = +0.21). This suggests that faster elongation driven by good codon-tRNA matching reduces ribosome stalling and co-translationally protects mRNAs from nuclease access, dynamically stabilising the most codon-optimised transcripts in the ribosome-depleted fraction during the acute starvation window(66–68). Simultaneously, the reduction in pU and m5C in heavy polysomes may reflect a modification remodelling of abundant translation-factor transcripts as ribosome demand shifts, with better-adapted sequences being preferentially retained in the translating pool. Clusters 3 and 4 thus propose mirror images of the same logic: in the glycosylation program, a *cis*-encoded initiation burden drives translational exclusion and passive poly(A) decay; in the translation machinery program, a *cis*-encoded elongation advantage drives dynamic stabilisation and active retention in the translating pool. These contrasting mechanistic hypotheses are summarised in **Figure 8B**.

The other functionally defined programs are consistent with this framework but with different emphases. In the ribosome-biogenesis program (cluster 1), better codon-optimised transcripts retain more poly(A) in the light-ribosome fraction (ρ = +0.21), consistent with preferential Pab1 association protecting them from deadenylation (**Figure 7D**). In the amino acid/catabolic program (cluster 5), codon optimality (ρ = +0.39) and Kozak strength (ρ = +0.22) both correlate with δSTE simultaneously, suggesting a rheostat-like regime in which the aggregate quality of *cis*-architecture across initiation and elongation steps scales proportionally with translational outcome, without any single step being rate-limiting. Neither cluster exhibits the sharp mechanistic decoupling seen in cluster 3, where Kozak context fails to predict outcome despite population-level weakness, reinforcing the view that cluster 3’s repression is specifically imposed by a multi-layered initiation barrier rather than tuned by any single *cis* parameter.

### 4.7. Limitations and broader implications

A broader implication concerns what happens to transcripts once translation is denied. The *cis*-encoded initiation barrier we infer for the glycosylation program predicts that these transcripts fail to load ribosomes and accumulate in the light-ribosome fraction. Independent work has established that glucose starvation in yeast rapidly sorts mRNAs by translational status: poorly-translated transcripts are sequestered into P-bodies and stress granules and can later return to polysomes when glucose is restored(69). Crucially, this sorting is sequence-directed, with transcript and promoter features determining whether an mRNA is translated and diffusely localised or excluded and concentrated into foci(70). Our feature-resolved, transcriptome-wide analysis is consistent with and extends this view: it is not a single determinant but the joint *cis*-architecture of a transcript with all, its initiation context, uORF burden, and structural accessibility, that appears to specify its translational fate, providing candidate molecular features that could underlie the sequence-directed sorting previously observed microscopically. Whether the cluster 3-transcripts defined here are physically enriched in starvation-induced granules is a direct and testable prediction of this framework.

The study is limited to a single, early (10-minute) starvation timepoint, capturing only the immediate post-transcriptional response before substantial transcriptional reprogramming has occurred; it does not address the dynamics of regulatory remodelling, and the extent to which the five-program architecture persists or dissolves at longer starvation times is unknown. DRS-based modification detection has lower per-site sensitivity than targeted approaches, and some modification sites, particularly those of low stoichiometry, will have been missed; the modification-related findings should be viewed as lower bounds on the extent of epitranscriptomic remodelling. The RBP regulon analysis is limited to RBPs with available yeast target data, and the RIP-chip-derived target sets vary in resolution and condition-specificity. Finally, while the lasso mixture model identifies that non-additive, cluster-conditional feature effects are statistically necessary to fit the data, this alone does not rule out the possibility that unmodelled confounders correlated with cluster membership account for some of the apparent interaction structure.

Importantly, the framework developed here, in which a single DRS experiment simultaneously measures multi-omic RNA layers which are integrated via a joint regression model that allows feature effects to be cluster-conditional, is applicable in principle to any biological system where multiple post-transcriptional regulatory layers can be profiled in parallel. The yeast glucose starvation response provides a tractable and well-characterised test case, but the methodological conclusion extends beyond it: post-transcriptional regulatory features should not be assumed to have fixed, universal relationships to translational output. A given molecular measurement can be a strong positive predictor, a strong negative predictor, or entirely uninformative depending on the program membership of the transcript being considered, and this context-dependence is invisible to any single-omic analysis or to multi-omic approaches that test each layer’s marginal relationship to the outcome independently. The convergence of *cis*-regulatory and post-transcriptional dynamic features within each program further suggests that the mechanistic architecture of translational control may be both more redundant and more program-specific than is typically assumed, a picture in which transcripts are not simply regulated by individual molecular events but are defined, at the sequence level, for particular regulatory fates that starvation then enacts.

## 2. Methods

### 2.1. Data processing and uni-layer post-transcriptional feature analysis

#### 2.1.1. Data sources

All sequencing data analysed in this study were generated in Horvath *et al.* (2024)(13) and are described in full therein. Briefly, the dataset consists of long-read direct RNA sequencing (DRS) libraries prepared from ribosome-disome-trisome (RDT) and higher polysome (PS) RNA fractions of *S. cerevisiae* (strain BY4741, an S288C derivative) sampled under non-starved conditions and after 10 minutes of glucose starvation. Libraries were sequenced on an Oxford Nanopore Technologies platform using the SQK-RNA002 chemistry. All raw sequencing data are publicly accessible at the NCBI Sequence Read Archive (SRA) under BioProject PRJNA1022817.

#### 2.1.2. Basecalling and Mapping

The raw FAST5 files direct RNA sequencing library were collected and basecalled into FASTQ using Guppy [version v5.0.7]. The FASTQ reads were further aligned with the reference transcriptome constructed as described(29) from SGD [version from 13.06.2024,(30). Alignment was performed using minimap2 [2.31-r1302,(31)] with option “ax map-ont -k14 -N10” and secondary alignments were filtered using samtools [1.23.1,(32)] command view with –F 2324 option.

#### 2.1.3. RNA Modification detection

RNA modifications (5-methylcytosine, m5C; N6-methyladenosine, m6A; and pseudouridine, pU) were identified at single-molecule resolution using SWARM [downloaded from github on September 2025;(33)], following the recommended pipeline for Oxford Nanopore Technologies (ONT) direct RNA sequencing with SQK-RNA002 chemistry.

SWARM operates in two stages. In the first stage (read-level classification, “model 1”), each individual read is scored for modification status at candidate sites; this model was applied separately to each biological replicate. In the second stage (site-level aggregation, “model 2”), read-level calls are pooled across replicates within a condition-and-fraction group to produce site-level modification probability estimates and stoichiometry predictions. Replicate merging was performed after confirming inter-replicate concordance and clean condition-based separation in principal component analysis (PCA) of modification profiles. The following groupings were used: non-starved PS (replicates B and C), non-starved RDT (replicates A and B), starved PS (replicates B and C), and starved RDT (replicates A, B and C), see replicate selection section below. SWARM model 2 outputs a binary site-level classification (modified/unmodified) together with a stoichiometry estimate at each covered site.

For each gene, a transcript-level modification rate was computed as the number of modified sites divided by the total number of detectable sites of the corresponding nucleotide within the gene body, defined as sites covered by at least 20 reads.

For each gene and each modification type, two 21-long binary vectors of modification status were extracted by assessing modification status at each of the 21 first positions starting at the start codon, and last 21 positions ending at the stop codon.

#### 2.1.4. RNA Modification comparisons

Differential stoichiometry between conditions and fractions was assessed using the generalised linear model (GLM) testing framework implemented in SWARM, applied to model 2 output. To focus on biologically meaningful changes, three pre-filters were applied: only sites detected as modified in at least one condition were considered (s=FDR_thresh); only sites with stoichiometry greater than 0.1 in at least one condition were retained (p=0.1); and only sites with an estimated inter-condition stoichiometry difference greater than 0.1 were tested (delta=0.1). Multiple testing correction used the Benjamini-Hochberg procedure(34) at a false discovery rate of 1%.

#### 2.1.5. Replicate selection

We performed some quality-based filtering of the replicates based on the following criteria: from the abundance matrix obtained with Salmon [1.10.1,(35)], we performed a PCA analysis on the samples and replicates which isolated replicate A of the non-starved PS and starved PS conditions as outliers. Additionally, NS_RDT replicate C stood out with abnormally low mapping rate (70 % compared to 97-98% for other samples), abnormally high modification rate and much lower per site stoichiometry correlation with other samples (0.825 on average) compared to in-between other samples correlations (0.987 at the lowest), a very strong indication of sample contamination. For all those reasons, we discarded replicates NS_PS_A, S10_PS_A and NS_RDT_C from downstream analysis.

#### 2.1.6. Poly(A) estimation and comparison

The poly(A) tail length from direct RNA sequencing was estimated using Nanopolish poly(A) analysis module [v10.2, (36)] from the raw fast5 files together with corresponding aligned bam files. Only reads with QC tag PASS were considered for further analysis. Violin plots of the mean poly(A) length within each sample was then plotted in R version 4.4.2, and comparisons between fractions were evaluated with a t-test rejecting the null hypothesis with Benjamini-Hochberg FDR correction 0.01.

#### 2.1.7. Differential mRNA degradation measurement

Transcript integrity was measured using INDEGRA [version 1.2.0 “Echidna”(37)] from in-house transcriptome aligned and filtered bam files processing each replicate separately, using the recommended pipeline. Biological degradation rates was extracted using the compareDegradation command treating all replicates as independent samples, and extracting the Tau_EAP estimates from each sample.

Comparisons between starved and non-starved conditions were performed on each of the polysomal fraction independently, with the compareDegradation command with option compareMaxNfl 100 using default prior p=0.1.

#### 2.1.8. Differential transcript abundance within polysomal fractions

To identify transcripts whose relative contribution to each polysomal fraction changes between non-starved (NS) and glucose-starved (S10) conditions, per-transcript read counts were extracted from processed alignment files and merged into a count matrix across all libraries; missing values were set to zero.

Differential abundance was analysed using DESeq2 [version 1.44(38)] with a combined condition-and-fraction factor as the design variable (design: ∼ConditionFraction, with levels NSPS, NSRDT, S10PS, S10RDT). This design permits independent contrasts to be extracted within each fraction without assuming a shared starvation effect across fractions. Genes were retained if at least five samples carried a read count of five or more; all others were excluded prior to model fitting. Size-factor estimation and dispersion modelling followed the DESeq2 standard pipeline.

Two contrasts were extracted independently: NS *versus* S10 within the polysome-associated (PS) fraction, and NS *versus* S10 within the ribosome-disome-trisome-associated (RDT) fraction. Transcripts were considered differentially represented if the Benjamini-Hochberg adjusted p-value was below 0.01 and the absolute log₂ fold-change exceeded 1.

#### 2.1.9. RNA structure predictions

Per-nucleotide RNA secondary-structure propensities were computed for every transcript in the annotated set. For each gene a mature mRNA sequence was assembled from the *Saccharomyces cerevisiae* S288C reference genome (assembly R64-4-1) by concatenating the annotated 5ʹUTR, coding sequence and 3ʹUTR, using the empirically defined UTR lengths. Each full-length transcript was folded with RNAfold from the ViennaRNA package [v2.6.4; (39)] in partition-function mode, disallowing isolated base pairs (--noLP) and with the folding temperature set to 30 °C to match the experimental growth conditions. From the resulting McCaskill base-pairing probability matrix we derived, for every position i, the probability that the nucleotide is base-paired, computed as the sum over all partners j of the pairing probabilities p(i,j) (the per-position unpaired probability is one minus this value); higher values indicate more strongly structured, less accessible positions.

These per-position pairing probabilities were summarised into transcript-level features in two complementary ways. First, for the joint model, each transcript region (5ʹUTR, CDS, 3ʹUTR) was summarised by its mean, three quartiles (Q1, median, Q3), variance, and the fraction of positions with a pairing probability above 0.9, yielding eighteen region-resolved structure features (six statistics × three regions); regions absent from a given transcript (for example genes without an annotated 5ʹUTR) were recorded as missing. Second, for the analysis of initiation-proximal structure, the mean pairing probability was taken over three windows defined relative to the main AUG (the entire 5ʹUTR, a start-codon window spanning the AUG ± 30 nt, and the first 50 nt of the CDS) and each cluster was compared with the union of all other clusters by a two-sided Mann-Whitney U test with the Cliff’s δ effect size(40), controlling the false-discovery rate across all cluster × window tests (Benjamini-Hochberg).

#### 2.1.10. Other features

Polysome Sedimentation Factor (PSF): was computed from DRS-based read counts in the PS and RDT fractions, following the procedure described in Horvath *et al.* (2024)(13). Per-transcript polysome abundance (PA) was calculated by first excluding transcripts with fewer than 20 cumulative reads across all conditions and fractions, then normalising non-zero counts by library size and imputing zero counts to 0.1. The PSF for each transcript was defined as PSF = PA_PS / (PA_PS + PA_RDT), providing a fractional measure of each transcript’s engagement with heavy polysomes relative to its total translated pool. To enable condition-wise comparisons, a differential δPSF was computed as δPSF_S10 = PSF_S10 / (PSF_S10 + PSF_NS), mirroring the self-normalising format used for δSTE and allowing direct quantitative comparison between the two metrics across the transcriptome

Gene sequence composition: for each gene, the transcript sequence was decomposed into three regions, 5ʹUTR, CDS and 3ʹUTR, using the custom transcriptome annotation described above. Within each region, sequence composition was summarized by region length and the fractional abundance of each nucleotide (A, T, G and C), yielding 15 sequence-based features per transcript. These features were included as covariates in the multi-layer integration model to account for potential sequence-composition biases in modification detection and degradation rate estimation.

### 2.2. Multi-layer post-transcriptional feature integration and cluster analysis

#### 2.2.1. Feature compilation

First, the dataset was assembled by integrating measurements from all regulatory layers characterised in this study. For each transcript, 236 features were compiled for non-starved (NS) and starved (S10) conditions, comprising: stochastic translation efficiency (STE) estimated with two different models referred as M1 and M2 (4 features), sequence composition (length and nucleotide frequencies of the 5ʹUTR, CDS and 3ʹUTR; 12 features), RNA modification stoichiometry and modification rate changes in both the polysome-associated (PS) and ribosome-dissociated (RDT) fractions for m5C, m6A and pseudouridine (24 features), differential poly(A) tail lengths in each fraction (4 features), differential degradation rates in each fraction (4 features), the polysome sedimentation factor and its stress-induced change (PSF; 2 features), quantile-based summaries of per-nucleotide structure assembly probability within each transcript region (18 features), positional modification features that indicates for m5C, m6A and pseudouridine in both PS and RDT fractions wheres there was a modication on the 21-start and 21-stop windows (168 features).

To investigate the molecular determinants of differential stochastic translation efficiency (δSTE) during glucose starvation, transcript-level differential features were computed as the difference between S10 and NS values. This resulted in 20 differential features, including δSTE estimated with both models (2 features), differential RNA modification stoichiometry and modification rate changes (12 features), differential poly(A) tail length (2 features), and differential degradation rates (2 features). Positional modification features were encoded using a discrete four-state scheme (0: no modification in either condition; 1: modification in NS only; 2: modification in S10 only; 3: modification in both conditions), reducing the original 168 positional features to 84 consolidated features. All remaining global transcript features were retained unchanged. After preprocessing and feature consolidation, the final dataset comprised 133 features: 84 positional modification features, 19 differential features, and 30 global features.

#### 2.2.2. Data imputation

Imputation was performed separately for the NS and S10 conditions to prevent cross-condition information leakage. Features that were shared between conditions (global features) were imputed using the NS dataset. Missing values were then imputed using an iterative machine learning approach based on Random Forest regression. For each feature containing missing values, a regression model was trained using the remaining observed features as predictors, and the missing entries were subsequently predicted. This procedure was repeated iteratively across all features for multiple passes, allowing progressively updated estimates to refine the imputations. For binary features, the regression outputs were converted into discrete values using a threshold of 0.5, where values ≤ 0.5 were assigned to 0 and values > 0.5 were assigned to 1. This ensures biologically interpretable binary states while preserving the continuous prediction strength of the model. Overall, this approach provides a robust, non-parametric imputation strategy that captures nonlinear relationships between genomic features, consistent with established Random Forest-based imputation methods(41).

#### 2.2.3. Autoencoder on modification windows

To reduce the dimensionality of the positional modification features, we reduced redundancy using a nonlinear dimensionality reduction approach. An autoencoder was trained to compress the 84-dimensional input space into a compact 2-dimensional latent representation. The model is based on a symmetric encoder-decoder neural network, where the encoder progressively compresses the input through several dense layers into a low-dimensional latent space, and the decoder reconstructs the original features from this representation. To optimise performance, we performed a randomised hyperparameter search over architectural parameters (layer sizes, activation functions, dropout rate, and regularisation strength), selecting models that minimised reconstruction error (mean squared error). Early stopping was used to prevent overfitting during training. After selecting the best hyperparameters, the final model was retrained on the full dataset. The resulting dataset comprised 51 features: the 2 latent dimensions learned by the autoencoder, 19 differential features, and 30 global features. The autoencoder was implemented using TensorFlow [Keras API(42)].

#### 2.2.4. Correlation with STE

We performed an association analysis to identify genomic features linked to STE-related phenotypes (STE_M1 and STE_M2). We systematically tested each feature for association with the STE variables. For each feature, we computed Pearson and Spearman correlation coefficients to capture linear and monotonic relationships, as well as the Maximal Information Coefficient (MIC, (43)) to detect potential non-linear dependencies. Features were considered associated based on statistical significance after multiple testing correction (Benjamini-Hochberg procedure, adjusted p-value < 0.01) and MIC values above the first quartile of the MIC distribution, ensuring retention of features with relatively strong non-linear associations. This combined approach enabled the identification of both linear and non-linear relationships between STE features and the remaining genomic variables. Based on these association criteria, 29 features (hereafter referred to as common features) were retained for the clustering step. The full set of 49 features was retained for downstream gradient boosting analysis, as that method handles non-linear relationship natively.

#### 2.2.5. Mixture of LASSO-penalised linear regression models

To identify groups of transcripts sharing a common linear regulatory architecture, a finite mixture of regression models was fitted to the transcriptome-wide feature matrix using the flexmix package [version 2.3.20; (44)] in R. The model assumes that transcripts are drawn from K latent subpopulations, within each of which δSTE is a linear function of the common features:

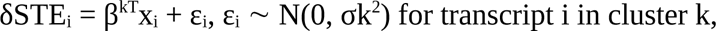

where x_i_ is the 29-dimensional vector of scaled common features, β^k^ is the cluster-specific coefficient vector, and σ^k^ is the cluster-specific residual standard deviation. Component membership was estimated by the expectation-maximisation (EM) algorithm. To promote sparsity and reduce the influence of collinear or weakly informative features, LASSO regularisation (α = 1) was applied to the component-specific regression coefficients within the M-step, using the FLXMRglmnet model specification in flexmix, which internally calls the glmnet package [version 5.0;(45)]. Prior to model fitting, all 29 common features were centred and scaled to unit variance across all transcripts; columns identified as co-linear by QR decomposition were excluded. Model fitting was performed independently for the two δSTE definitions (M1 and M2) and independently for regular (unpenalised; FLXMRglm) and LASSO-penalised (FLXMRglmnet) component specifications. The number of components K was varied from 1 to 6; the K = 1 model, equivalent to a single global regression, served as a performance baseline. All fits used a fixed random seed (123) to ensure reproducibility.

#### 2.2.6. Cluster number selection

The optimal number of mixture components K was selected using two information-theoretic criteria computed from the fitted model objects. The Bayesian Information Criterion [BIC(46)] penalises the log-likelihood by model complexity:

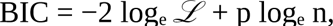

where ℒ is the maximised likelihood, p is the number of free parameters, and n is the number of transcripts. The number of mixture components with minimal BIC was retained for downstream analysis. As a complementary elbow-plot diagnostic, the weighted within-cluster MSE of the fitted mixture regression was also computed: for each K, predicted δSTE values were obtained by applying the cluster-specific LASSO coefficient vectors to the scaled feature matrices of the respective cluster members, and the MSE was normalised by the global variance of δSTE and averaged across clusters weighted by cluster size.

#### 2.2.7. Gradient boosting and SHAP-based feature importance

To characterise the predictive contribution of all 49 RNA features to δSTE within each cluster identified by the LASSO mixture model, a gradient boosted tree model was fitted independently per cluster using the xgboost package (version 1.7.11.1(47)). Unlike the mixture regression, gradient boosting captures non-linear effects and feature interactions, and it is not restricted to the 29 common features: the full 49-feature matrix was supplied as input, with missing values handled natively by the tree-splitting algorithm. Unbiased performance estimates were obtained via a nested cross-validation scheme with five outer and five inner folds. In each inner loop, hyperparameters were selected by grid search over the following values: number of boosting rounds ∈ {100, 300}, maximum tree depth ∈ {3, 6}, learning rate ∈ {0.05, 0.1}, column subsampling fraction ∈ {0.6, 0.8}, and row subsampling fraction ∈ {0.7, 1.0}, with minimum child weight and γ fixed at 1 and 0 respectively. Model performance was summarised as the mean squared error (MSE) and mean absolute error (MAE) on held-out outer folds, normalised by the range of δSTE values in the corresponding training set.

Feature contributions were quantified using TreeSHAP(48), the exact Shapley-value decomposition for tree ensembles, obtained via the predcontrib argument in xgboost. For transcript i and feature j, the SHAP value φᵢⱼ represents the contribution of feature j to the predicted δSTE relative to the global mean, with the additive constraint that Σⱼ φᵢⱼ equals the deviation of prediction i from the mean. Feature importance within each cluster was summarised as the mean absolute SHAP value across all cluster members (mean|φⱼ|). The directional effect of each feature was characterised by the sign of the Pearson correlation between feature values and the corresponding SHAP values within the cluster: a positive correlation indicates that transcripts with a higher value of the feature tend to receive a positive SHAP contribution, *i.e.* the feature pushes δSTE upward in that cluster, consistent with the sign convention used for the LASSO regression coefficients.

#### 2.2.8. Gene Ontology enrichment analysis

To assess the biological coherence of the identified transcript clusters, Gene Ontology (GO) enrichment analysis was performed for the Biological Process (GO:BP) sub-ontology using the clusterProfiler package [version 4.12.6(49)] in R, with the *Saccharomyces cerevisiae* annotation database org.Sc.sgd.db. ORF identifiers were mapped to Entrez gene identifiers using the bitr function. For each cluster, a hypergeometric over-representation test was applied against the background of all 5,033 analysed transcripts, with multiple testing correction by the Benjamini–Hochberg procedure at a false discovery rate threshold of 0.05. Cross-cluster comparison was performed using the compareCluster function, which applies the enrichment test simultaneously to all cluster gene lists and produces a unified dot plot in which dot size encodes the gene ratio (proportion of cluster genes annotated to the term) and dot colour encodes the adjusted p-value. Hierarchically redundant GO terms were not collapsed algorithmically; only the most specific non-redundant terms are discussed in the main text.

#### 2.2.9. UTR and CDS sequence extraction

Annotated 5ʹUTR, CDS, and 3ʹUTR boundaries were defined as described earlier(29). For each transcript the three regions were extracted from the reference FASTA file: the 5ʹUTR as the first L_5 nucleotides, the 3ʹUTR as the last L_3 nucleotides, and the CDS as the intervening sequence, where L_5 and L_3 are the annotated UTR lengths. Genes were required to have an annotated UTR length of at least 10 nt for UTR analyses; for CDS analyses a minimum coding sequence length of 100 nt was required. One gene for which the annotated UTR length exceeded the deposited sequence length was excluded throughout.

#### 2.2.10. K-mer Shannon entropy

For each gene and each region, k-mer Shannon entropy was computed for k = 1 to 6. Overlapping k-mers of length k were enumerated with a sliding window (step = 1), yielding L − k + 1 k-mers per sequence of length L. Shannon entropy was calculated as H = −Σ p_i_ log_2_(p_i_), where p_i_ is the observed proportion of k-mer i among all valid k-mers. The primary normalised metric was H_theo = H / (k · log_2_(4)), which scales entropy by the theoretical maximum for a uniform distribution over 4^k^ k-mers and ranges from 0 (one k-mer type) to 1 (uniform distribution). Only genes with a UTR longer than max(10, 4^k^) nt were retained for analysis for each k value. To test whether differences in k-mer entropy between clusters were independent of UTR length, we fitted an ANCOVA model H_theo ∼ log_10_(UTR_length) + cluster for each UTR type and k-mer order (k=1-6). The significance of the cluster term (beyond the length covariate) was assessed via an F-test comparing nested models (with *vs.* without cluster), with p-values BH-corrected across all 12 tests (6 values of k × 2 UTR types). Effect sizes are reported as partial eta-squared (η^2^_partial), defined as SS_cluster / (SS_cluster + SS_residual), which quantifies the proportion of residual variance explained by cluster membership after accounting for UTR length.

#### 2.2.11. Nucleotide composition analysis

For each transcript region, the fractional abundance of each nucleotide (A%, T%, G%, C%) and the combined AU content (A+T%) were computed directly from the extracted sequence. Between-cluster differences in nucleotide composition were assessed with the same ANCOVA framework described above (composition ∼ log_10_(region length) + cluster) applying an F-test for the cluster term and BH correction across the nucleotides tested within each region. For the CDS, the pre-computed nucleotide percentages from the transcript feature matrix (A_percent_CDS, C_percent_CDS, T_percent_CDS) were used in place of direct sequence extraction. Length-adjusted pairwise comparisons used estimated marginal means (emmeans, BH correction).

#### 2.2.12. CDS codon usage enrichment analysis

To test whether between-cluster differences in CDS k-mer entropy reflect genuine codon usage differences rather than non-specific nucleotide composition effects, k-mer enrichment was assessed for clusters 3 and 4 against all remaining clusters combined, using two complementary approaches matched to the biological interpretability of each k value. For k = 3 and k = 6, a frame-aware analysis was applied: only k-mers whose start position falls at a codon boundary (positions 1, 4, 7, … in the CDS; *i.e.*, positions ≡ 1 mod 3 in 1-based indexing) were counted. This restricts the analysis to in-frame codons (k = 3) and consecutive codon pairs (k = 6), and permits direct comparison with known codon usage tables without confounding contributions from overlapping reading frames. k-mer counts were aggregated across all sequences within the focus cluster and across all sequences in the remaining clusters. For each k-mer, enrichment in the focus cluster was assessed with a two-sided Fisher’s exact test on the 2 × 2 contingency table of [k-mer count in focus cluster, remaining k-mer count in focus cluster] *versus* [k-mer count in other clusters, remaining k-mer count in other clusters]. k-mers with a total count below 10 across both groups were excluded. The log₂ fold-change was computed as log₂((f_focus + ε) / (f_rest + ε)), where f denotes k-mer frequency (count / total k-mers in the group) and ε = 10^−9^ is a pseudocount. P-values were corrected within each k using the Benjamini–Hochberg procedure. Frame-aware k-mer counting was implemented with a custom R function; sliding-window counts used the oligonucleotideFrequency function from Biostrings [version 2.72.1;(50)].

#### 2.2.13. Codon optimality indices

The Codon Adaptation Index (CAI;(51)) was computed for each transcript’s CDS using the 130 cytoplasmic ribosomal-protein genes (RPL*/RPS*) as the high-expression reference set, with relative synonymous codon usage (RSCU) estimated from pooled reference codon counts (Laplace pseudocount 0.5) and per-gene CAI as the geometric mean of relative adaptiveness w over all scored codons (initiator AUG, stop codon, and single-codon amino acids Met and Trp excluded). The tRNA Adaptation Index (tAI;(52)) was computed in parallel using tRNA gene copy numbers from the GtRNAdb sacCer3 set (275 nuclear genes; the elongator-decoding vector excludes the initiator tRNA-iMet, leaving 268 genes over 41 sense anticodons), with optimised eukaryote wobble penalties (U:G 0.41, U:C 0.28, A:G 0.9999, C:G 0.68; all other pairs 0). Zero-adaptiveness codons were set to the geometric mean across all codons; per-gene tAI is the geometric mean of w excluding initiator AUG and stop codons. Both indices were verified against expected yeast biology (genome-wide CAI–tAI Spearman ρ = 0.93, consistent with the canonical value of ∼0.9; expected optimal/non-optimal codon ranking recovered). Per-cluster differences *versus* background (all other clusters) were assessed with the Mann-Whitney U test and reported as Cliff’s delta(40); multiple testing was controlled by BH-FDR within each analysis.

#### 2.2.14. Initiation context, uORFs, and RNA secondary structure

Extended Kozak context was scored for each transcript as the position-weight match of the −6 to +4 window around the initiator AUG against a universe position-frequency matrix(53). uORFs were identified as AUG-to-in-frame-stop reading frames within each annotated 5ʹUTR and summarised per transcript as uORF prevalence (binary: any uORF present) and uAUG density (upstream AUG count per 100 nt of 5ʹUTR; B3). RNA secondary structure was estimated from per-nucleotide base-pair probabilities computed with RNAfold (ViennaRNA package(39)), sliced into three windows per transcript: the full 5ʹUTR, a ±30 nt window centred on the initiator AUG, and the first 150 nt of the CDS. Each window was summarised as its median P(paired), and a positional metagene profile was constructed over offsets −100 to +150 around the AUG. Sequence-specific RBP binding motifs (Puf3/4, Pub1/ARE U-rich, Whi3, Ssd1, Vts1/SRE, Khd1) were scanned as IUPAC patterns in each transcript region. All per-cluster contrasts used all other clusters as background and were assessed with Cliff’s delta and BH-FDR.

#### 2.2.15. Experimental RBP target enrichment

Curated RBP regulon sets were derived from the Hogan *et al.* (2008)(54) genome-wide RIP-chip atlas: from the SAM-FDR matrix (Dataset S2), a transcript was assigned to an RBP regulon when any probe reached SAM FDR ≤ 1% (probes with negative FDR values, indicating SAM under-enrichment, were excluded); probes were collapsed to systematic ORF identifiers, yielding 42 non-empty target sets covering 7,788 gene-RBP assignments (four atlas RBPs had no targets at this threshold). Puf1-5 sets were independently cross-validated against the Gerber *et al.* (2004)(55) RIP-chip tables (Hogan-vs-Gerber Jaccard 0.42–0.53 for Puf2-5). For each cluster and each of the 42 RBP sets, a two-sided Fisher exact test with a Haldane-Anscombe-corrected log_2_ odds ratio was applied(56,57), comparing cluster membership against all other clusters. BH-FDR was computed across all 168 cluster × RBP tests simultaneously.

#### 2.2.16. Cross-cluster synthesis matrix

Effect sizes from all feature analyses were assembled into a single oriented signal matrix with one row per feature family and one column per cluster. Each signal was sign-oriented to a common pro-translation axis (positive = feature configuration favouring higher δSTE): δSTE (median), CAI (Cliff’s δ *vs.* background), tAI (Cliff’s δ), Kozak strength (Cliff’s δ), uORF prevalence (−log_2_ odds ratio, negated so that uORF depletion = pro-translation), start-codon accessibility (−Cliff’s δ of P(paired), so lower pairing = positive), and Pab1-target enrichment (log_2_ OR from m07). Two coupling signals, the partial Spearman ρ(Kozak, δSTE | length) from C2 and Cliff’s δ of uORF repression from C3, are shown verbatim. Figure cells are coloured by per-row z-score across the five clusters to highlight relative pattern; native effect sizes and significance stars are annotated within cells.

## Acknowledgements

The authors are grateful to the Oxford Nanopore Technologies for helpful and insightful discussions and collaborative spirit, with special thanks to Warren Bach, Mike Yarski, Steven Batinovic, Nao Goto, Sam Dyer, Angela Von and Ross Napoli. The authors would like to acknowledge their use of the Kaya HPC facility at UWA and the unwavering, highly enabling support provided by the Kaya HPC team, including the amazing contributions and work of Chris Bording.

## Author Contributions Statement

N.S. and A.C. conceived, funded, managed and supervised the work, all authors performed formal analyses and coding, figure generation and data interpretation, O.R. produced the original manuscript draft with the input of N.S. and A.C., O.R., N.S, and A.C. wrote and edited the final manuscript with the input of S.M.

